# A deep-learning–informed prior and Bayesian model for differential AP-MS interactome analysis

**DOI:** 10.64898/2026.07.06.736690

**Authors:** Manuel Seefelder

**Author notes:** **Corresponding author:** Manuel Seefelder, Department of Gene Therapy, Ulm University, Helmholtzstraße 8/1, D-89081 Ulm, Germany.

## Abstract

Affinity-purification mass spectrometry (AP-MS) maps a bait protein’s partners, but every purification also captures abundant non-specific background that masks genuine interactors. Established tools such as SAINTexpress and CompPASS score one evidence type, treat correlated signals as independent, and ignore prior knowledge of likely interactions. BayesInteractomics, an open-source Julia framework, addresses both limitations by combining machine learning with Bayesian statistics. A neural network trained on protein structures predicts direct binding. A calibrated meta-learner turns this into an informed prior. The prior guides a Bayesian copula-mixture model integrating three AP-MS evidence streams: enrichment, co-abundance, and detection reproducibility. Each candidate receives an interaction probability at a controlled false-discovery rate, optionally updated by structural docking. On synthetic data it ranks first in every benchmark (median AUROC 0.747), and across independent studies it raises high-confidence-call reproducibility from 21% to 79%. It also identifies which interactions are gained or lost between two conditions, unlike established tools.

## Main

The intricate network of protein-protein interactions (PPIs) forms the backbone of cellular machinery, governing nearly every biological process from signal transduction to metabolic regulation^12^. A primary technology for elucidating these networks is affinity purification mass spectrometry (AP-MS)^34^, which identifies protein complexes on a proteome-wide scale. Its key challenge is the robust distinction of *bona fide* interactors from a large background of non-specific binders and contaminants. Traditional analyses rely on simplistic statistics such as fold-change and p-values, which fail to capture the multi-faceted nature of the evidence and yield high rates of false positives and false negatives^5^; they also lack a principled way to integrate prior biological knowledge or to handle the experimental noise inherent in datasets pooled across diverse protocols and publications.

State-of-the-art methods such as SAINT and its derivatives advanced beyond fold-change by modelling prey-count distributions with a negative binomial to provide a probability score per putative interaction^6^. While powerful, they focus on a single evidence stream — prey enrichment from spectral counts — with no native framework for integrating other partially dependent evidence types such as bait-prey correlation or detection frequency, and no mechanism for incorporating data-driven prior knowledge from orthogonal sources such as *in silico* interaction predictions. A comprehensive framework combining the predictive power of modern deep learning on protein sequence and network data with a fully Bayesian evidence-combination model — yielding an actual posterior probability per interaction, and hence a reliable ranked candidate list for functional validation — remains an open challenge.

Here we present *BayesInteractomics*, an open-source Julia framework for rigorous probabilistic analysis of interactomics data, asking whether a hybrid model combining a deep-learning–informed prior with a multi-evidence Bayesian mixture model can increase the confidence and specificity of interactome mapping. The pipeline introduces a Deep neural network (DNN) predicting direct protein-protein interactions and a copula mixture model that learns the dependency structure among three evidence streams — protein enrichment, bait-prey correlation, and detection frequency — estimated robustly by a maximum a posteriori expectation-maximisation (MAP-EM) algorithm with an empirical null.

We validate the approach on three complementary fronts. On synthetic data with complete, known ground truth we measure interaction accuracy and probability calibration directly, where comparators cannot be benchmarked against an incomplete reference. We benchmark head-to-head against SAINTexpress, CompPASS and MiST on two independent human AP-MS deposits (OpenCell and Hein *et al*.), scoring every tool against a curated CORUM ∪ IntAct reference — the closest approximation to ground truth on real data, against which an open-world, partly circular reference can corroborate but not crown a single most-accurate scorer. And through a meta-analysis of the huntingtin (HTT) interactome we demonstrate calibrated, differential interactomics, pooling independent studies of wild-type and polyglutamine (polyQ)-expanded HTT the detailed biological findings of that meta-analysis, and of the obligate HTT–huntingtin-associated protein 40 (HAP40) complex, are reported in dedicated companion studies^78^.

## Results

### A protein-protein interaction prediction model

#### Deep neural network

At the base of BayesInteractomics (Figure 1) is a DNN trained to predict direct physical contact from protein-sequence and cross-species STRING-network embeddings, with ground truth from PDB structures (a pair labelled a direct interactor when its minimum inter-chain *C*_*α*_ distance falls below 5Å). On a similarity-minimised held-out test set of *n* = 1,862 protein pairs disjoint from the 70/20/10 training split, the network selected by a 1,100-configuration random search discriminates direct from non-direct contacts with a ROC-AUC of 0.829, an Matthews correlation coefficient (MCC) of 0.517, and an area under the precision–recall curve (AUPRC) of 0.737 (Table 1, DNN-baseline row).

**Table 1:**
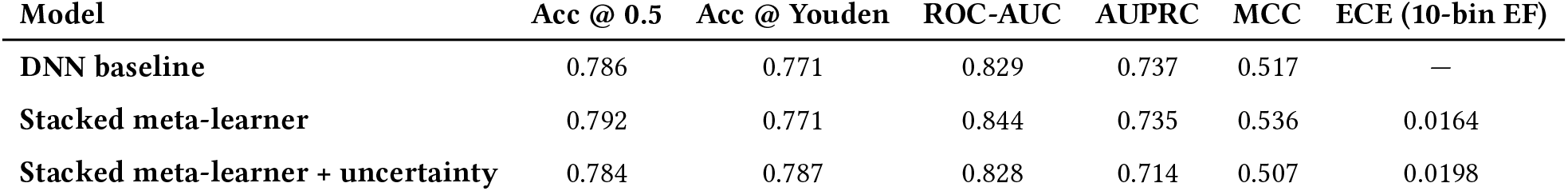
Held-out discrimination and calibration of the prior and the stacked meta-learner. Test-set performance on *n* = 1,862 protein pairs. The first meta-learner variant blends the 14 raw input features (in-species STRING, transferred STRING, protein-domain, and DNN prior); the “+ uncertainty” variant adds the Monte-Carlo-Dropout standard deviation from the DNN as a fifteenth column. Accuracy is reported at both the natural 0.5 probability threshold and the Youden-J-optimal threshold. The ECE column reports the 10-bin equal-frequency flavour used as the calibration acceptance metric; the DNN-only baseline cell shows “—” because its uncalibrated outputs are not subjected to that threshold.

**Figure 1.**
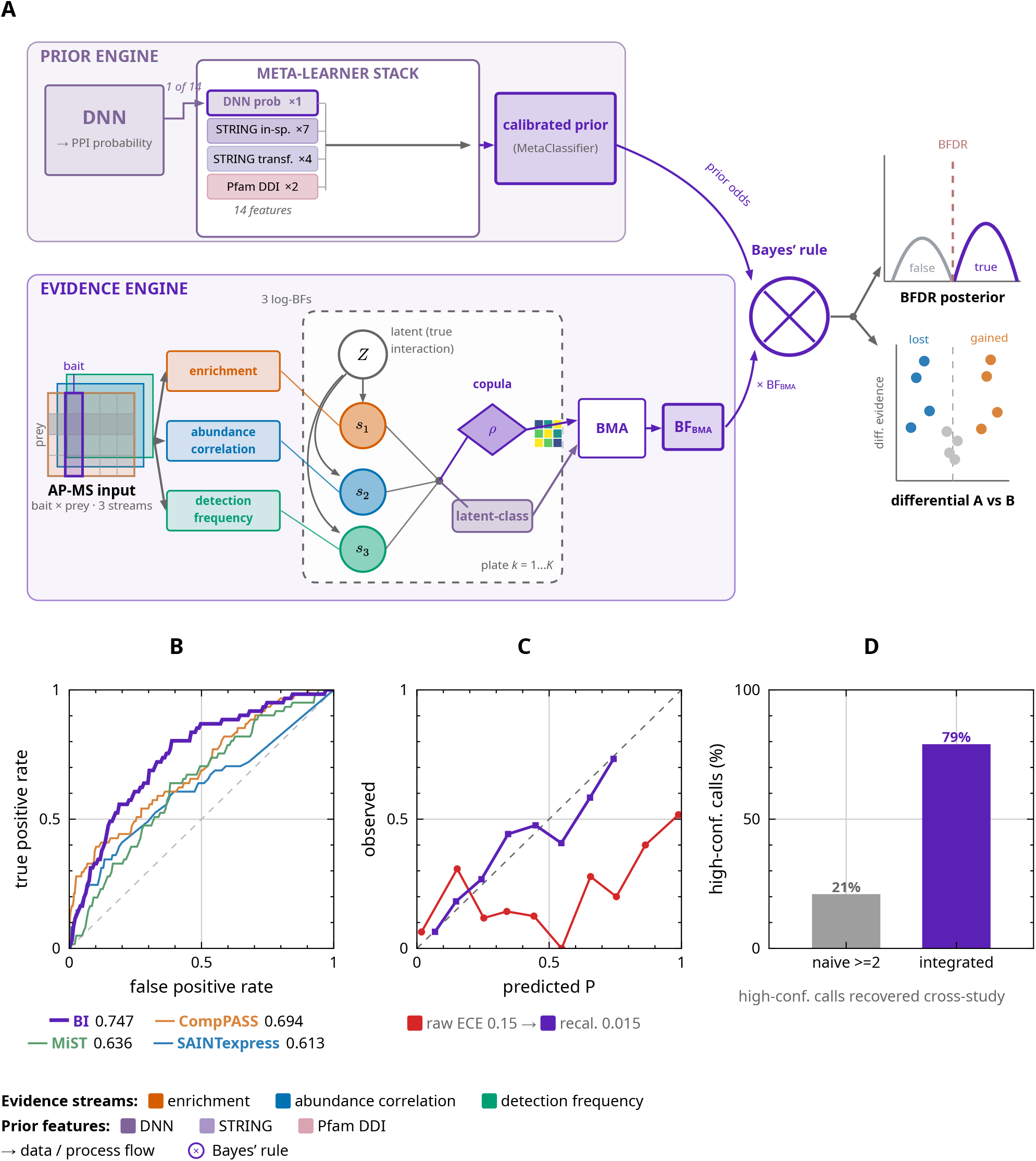
BayesInteractomics couples a sequence-and-database prior with a multi-evidence Bayesian model. (A) Two engines meet only at a Bayes’-rule node. The *prior engine* (sequence and database features only, no AP-MS data) treats a deep neural network’s interaction probability — trained on Protein Data Bank 3D contacts from sequence and STRING-network embeddings — as one of fourteen features (seven in-species STRING channels, four cross-species transferred STRING channels and two Pfam domain–domain features) fed to a stacked meta-learner (six base learners, an L2-regularised logistic blender and an isotonic calibrator); its calibrated output is the prior. The *evidence engine* (experimental AP-MS data) scores three streams — enrichment, abundance correlation and detection frequency — as log-Bayes-factors, fused through a copula mixture and a latent-class mixture and combined by Bayesian model averaging into BF_BMA_; the two engines combine as posterior odds = prior odds × BF_BMA_, controlled at a Bayesian false-discovery rate, with a native across-condition differential. Panel A is schematic. (B) Discrimination on the realistically-calibrated synthetic benchmark (ROC; BayesInteractomics first of four tools; full curves and paired statistics in Extended Data Fig. 4). (C) Posterior calibration, raw versus recalibrated (Extended Data Fig. 3). (D) Cross-study reproducibility of high-confidence calls, naive consensus versus the integrated model.

This held-out result is not a fortunate split: a 5-fold cross-validation re-training the identical architecture reproduces the same discrimination tightly (test ROC-AUC standard deviation 0.007 across folds), and a 5-model deep ensemble reaches a test ROC-AUC of 0.821, so the selected network sits inside the ensemble’s robustness band. The network also exposes a per-pair predictive uncertainty from *K* = 30 Monte-Carlo-dropout passes (*ρ* = 0.51 against the deep-ensemble disagreement), an optional feature for the meta-learner below.

We use this network to incorporate sequence- and data-base-derived information as prior information into our analysis pipeline. The network scores structural plausibility from sequence and network context alone, with no access to the experimental AP-MS evidence. The output of this network serves as one input to the meta-learner described below, whose calibrated prediction forms the prior the model then weighs against the experimental evidence.

#### Meta-learner

To integrate the DNN prior with the STRING evidence channels and transferred network features, we replace the earlier classifier sweep with a single stacked meta-learner: an `MLJ.Stack` of six fixed-hyperparameter base learners (histogram gradient boosting, EvoTrees, L2-regularised logistic regression, *k*-nearest neighbours, random forests, extra trees) blended by an L2-regularised logistic-regression layer trained on out-of-fold predictions and calibrated post-hoc with an isotonic transform. The schema and hyperparameters were selected under 5-fold group-disjoint cross-validation.

The meta-learner consumes 14 raw features: seven in-species STRING channels, four transferred STRING channels, two Pfam protein-domain features, and the DNN prior probability. The transferred-STRING and domain features are generated per species from each organism’s own STRING and Pfam resources, so a non-human bait is scored on species-correct evidence and the calibrated prior is species-general. Relative to the bare DNN baseline, stacking these channels and recalibrating lifts discrimination (ROC-AUC 0.829 → 0.844, MCC 0.517 → 0.536) while tightening calibration to a near-ideal expected calibration error (ECE) of 0.016 (Table 1). A second variant adding a fifteenth Monte-Carlo-Dropout-uncertainty column improves neither discrimination nor calibration, so we retain the 14-feature variant as the production default; both sit below the 0.035 calibrated-ECE threshold for downstream use. The blender’s calibrated probability is the only quantity the Bayesian layer consumes as a prior. Per-feature contributions and the feature-addition trajectory are shown in Extended Data Fig. 1 and Extended Data Fig. 2, respectively. Among the meta-learner’s 14 input features — as opposed to the four out-of-fold base-learner predictions the stack adds internally — STRING text-mining is the largest direct contributor (Extended Data Fig. 1 A).

### Discrimination and probability calibration

A probabilistic caller is only as trustworthy as the meaning of its numbers, so we interrogated the posterior on a graded-signal synthetic substrate with complete ground truth, each true interactor assigned an enrichment drawn uniformly over a wide range so the posterior spans the whole [0, 1] axis. On this substrate (*n* = 1500 candidates, 16.3% true interactors) the posterior *ranks* interactors strongly (area under the ROC curve 0.83) but its *raw* probabilities are over-confident, with an ECE of 0.15 and an maximum calibration error (MCE) of 0.55 (Brier 0.16): discriminative, but not yet a literal probability.

This over-confidence is a monotone distortion of the scale, not a defect of the ranking. A held-out five-fold Platt map fitted *against the known labels* collapses ECE 0.15 → 0.015 and MCE 0.55 → 0.14 and brings the reliability curve onto the diagonal (Extended Data Fig. 3), with isotonic regression reaching the same ECE (0.019); because the map is monotone the ranking is invariant (ROC area unchanged at 0.83). Given ground-truth labels the posterior is therefore near-perfectly calibratable.

### Synthetic benchmarking on a realistically-calibrated substrate

Because real references are incomplete and partly circular (as discussed below), scorer accuracy can be adjudicated directly against known truth only on synthetic data — and only if the substrate is realistic. Conventional spike-in simulators make the spectral-count branch almost perfectly separable, *inverting* the branch ordering seen in real data and handing the count-based comparators an artificial advantage (Extended Data Table 1). We therefore calibrated the generator’s per-prey discriminability to a real human deposit (OpenCell, PXD024909), tuning its negative-binomial count and Normal label-free quantification (LFQ) branches to the separability measured on real data — an external target fixed before scoring and not chosen to favour BayesInteractomics (Extended Data Table 1). BayesInteractomics was run copula-only — without its prior, which can only be computed for real proteins — so this result likely represents a lower bound on the accuracy of the production pipeline.

The calibration changes the benchmark decisively. The count-based comparators, near-perfect on idealised simulators (AUROC 0.97–0.99), collapse to a realistic 0.61 –0.69 (Extended Data Fig. 4A): their apparent synthetic dominance is a substrate artefact, not superior accuracy. Against this corrected baseline BayesInteractomics is competitive with or ahead of every comparator: across *B* = 7 independently seeded batches its median AUROC is 0.747 (versus CompPASS 0.687, MiST 0.636, SAINT-express 0.624), first in every batch, with each paired AUROC difference’s 95% bootstrap confidence interval entirely above zero (+0.134 over SAINTexpress, +0.068 over CompPASS, +0.128 over MiST; Wilcoxon signed-rank *p* = 0.016); on precision–recall it beats SAINTexpress (+0.083) and MiST (+0.112) and trails only CompPASS (−0.040; Extended Data Fig. 4B). Two caveats temper this comparison: the generator was calibrated to an external target, and BayesInteractomics ran without its prior. We therefore read the synthetic benchmark not as our primary reliability claim, but as evidence on two points — that the framework is not disadvantaged where ground truth is observable, and that conventional simulators can rank methods in the wrong order.

### Comparison with established tools on real AP-MS data

The synthetic results establish strong, rank-invariant discrimination; the more difficult question is how the full pipeline fares against existing tools on real data, where ground truth is unobtainable. We benchmarked it against SAINTexpress, CompPASS and MiST on a human AP-MS deposit (OpenCell, PXD024909; 76 baits, 2,377 reference positives among ∼104,000 candidate pairs), scoring every tool against an external CORUM ∪ IntAct reference (Figure 2). On the raw scores no method dominates: discrimination is modest and tightly clustered (ROC-AUC MiST 0.69, SAINTexpress 0.67, BayesInteractomics 0.66, CompPASS 0.61) and precision–recall area is low for every tool (AUPRC ≤ 0.12 against a 0.02 positive rate; Figure 2A–B).

**Figure 2.**
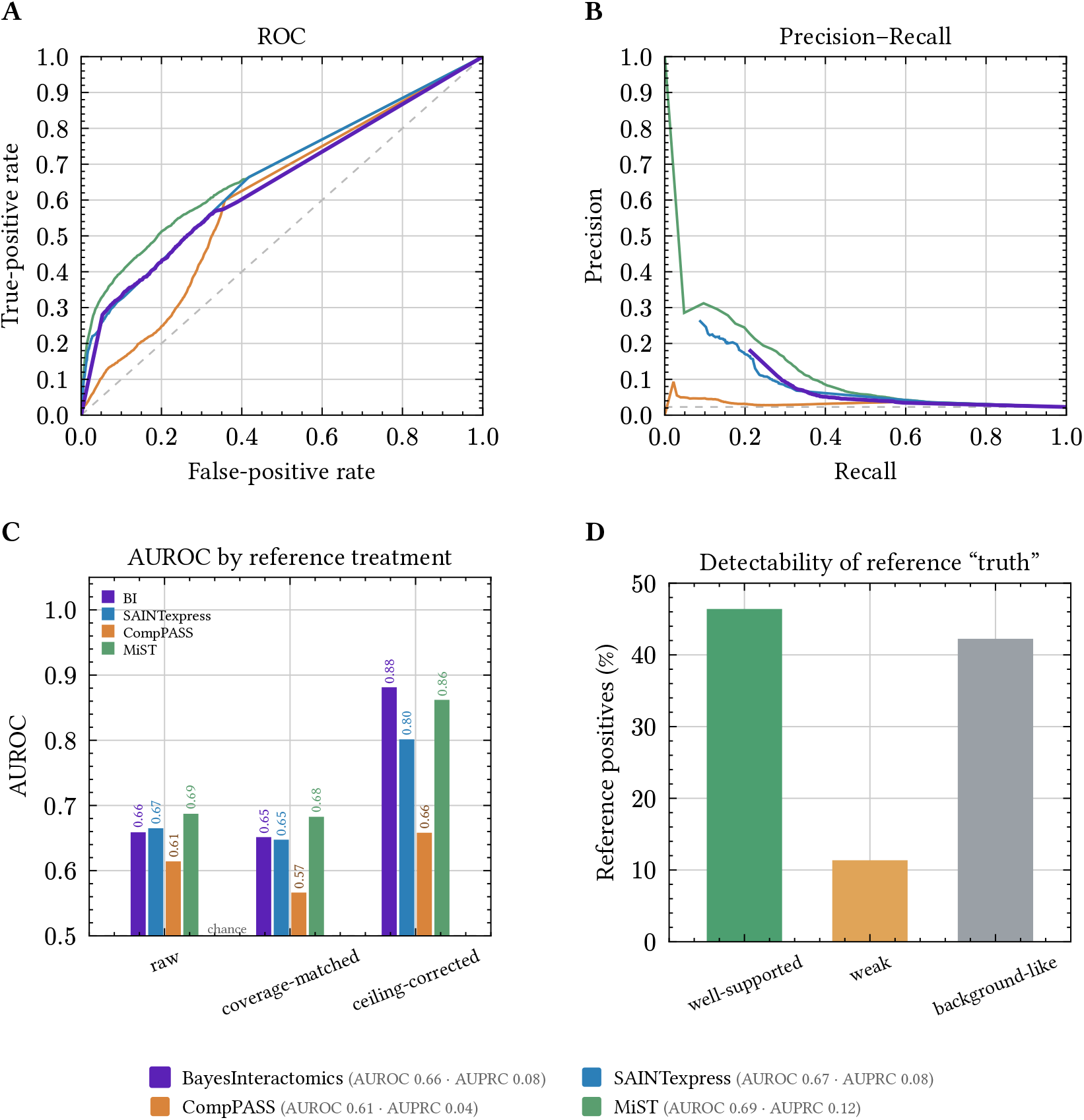
Real-data benchmark against established tools. All four scorers were run on the OpenCell AP-MS panel (PXD024909; 76 baits, 2,377 reference positives among ∼104,000 bait–prey candidates) and scored against an external CORUM ∪ IntAct reference assembled independently of any method; SAINTexpress and BayesInteractomics run per bait, CompPASS and MiST whole-panel. (A) Pooled ROC and (B) precision–recall curves on the raw scores; dashed lines mark chance (the diagonal and the class prior 0.02, respectively). (C) AUROC under three reference treatments — *raw* (every unscored candidate ranked last),*coverage-matched* (the common candidate set BayesInteractomics scores), and *ceiling-corrected* (positives restricted to the well-supported tier); bars start at the chance baseline (AUROC 0.5, dashed). (D) Detectability of the curated “truth”: of the reference positives, 46% are well-supported (≥ 1 log_2_ enrichment and detected), 11% weak, and 42% background-like (< 0.5 log_2_ enrichment). The reference is open-world (absence ≠ non-interaction), partly circular (seeded by SAINT/CompPASS-class pipelines), and recall-capped.

This raw ranking cannot adjudicate accuracy, for a reason we can measure on the data itself. Of the 2,377 curated “true” positives only 46% carry recoverable enrichment signal here (well-supported: ≥ 1 log_2_ enrichment and detected in the assay); a further 11% are weak (0.5 –1 log_2_ enrichment) and 42% are background-like (< 0.5 log_2_ enrichment), tiers no scorer can recover and that are therefore forced false negatives for every method (Figure 2D). This recall ceiling, the open-world precision cap and the partial circularity of the reference tilt the raw ranking toward the comparators. Under honest like-for-like comparisons the picture changes: coverage-matched to the candidates it scores, BayesInteractomics ranks second behind MiST, and restricted to the positives the assay can genuinely support it rises to match the strongest comparator and pull clear of the rest: its ROC-AUC of 0.88 is within sampling noise of MiST’s 0.86 but clear of SAINTexpress’s 0.80 and CompPASS’s 0.66 (Figure 2C), its posterior tracking per-prey enrichment as faithfully as the best comparator (Spearman *ρ* = 0.66 versus 0.65, 0.41, 0.16). We therefore read this benchmark as corroboration rather than as a ranking: against an incomplete, partly circular reference no AP-MS scorer can be decisively shown more accurate than another.

The same pattern reproduces on a second, independent human dataset in a contrasting overexpression regime (Hein 2015, 75 baits; Extended Data Fig. 5): raw and coverage-matched rankings are again reference-dominated, while on the well-supported positives BayesInteractomics again reaches the top tier: its ROC-AUC of 0.92 is within sampling noise of MiST’s 0.93 and clear of SAINTexpress’s 0.86 and CompPASS’s 0.85. An architectural ablation (Extended Data Fig. 6) is consistent: on synthetic truth the three evidence streams are mutually redundant for ranking, while on real data removing any single stream leaves discrimination largely intact but degrades calibration and threshold-call quality, with the enrichment stream contributing most.

### Structural docking of high-confidence interactors

The optional structural-docking step folds a structural Bayes factor into the posterior, derived from an interface-quality score benchmarked against complex predictions of known quality. On the published AlphaFold2/ AlphaFold3 complex-scoring benchmark^9^, a four-metric re-derivation of the AlphaFold3-calibrated interface composite (interface pLDDT, interface PAE, pTM and ipTM) separates correct from incorrect predicted complexes at an area under the ROC curve of 0.93, against 0.83 for the widely used pDockQ. Adding the score’s fifth term, a VoroIF graph-neural-network feature, did not improve discrimination (0.93 → 0.92), so we retain the parsimonious four-metric form. Its application to real interactome biology is demonstrated in the companion HAP40 study^8^ below.

### Meta-analysis of the HTT-interactome

Unlike SAINTexpress, CompPASS and MiST — each scoring interactors against control within a single condition — BayesInteractomics models the across-condition comparison directly, contrasting the full per-prey evidence in the Bayes-factor domain (Equation 25), propagating per-condition uncertainty into a calibrated differential call, and ranking proteins by decision-theoretic validation priority. A companion meta-analysis exercises this on the HTT interactome^7^, pooling four published AP-MS datasets^10111213^ spanning mouse and human tissues with distinct tags, epitopes and polyQ lengths into one calibrated model placing wild-type and polyQ-expanded baits on a common posterior scale. Of 4,338 proteins evaluated, 275 were condition-dependent — 70 lost on expansion, 205 gained — atop a 157-protein constitutive core both baits engaged equally, including the obligate partner HAP40 (`F8A2`) and the TRiC/CCT chaperonin, a polyQ-independent positive control recovered across all four datasets. The analysis showcases three capabilities per-study fold-change or spectral-count scores cannot provide (Figure 3). First, differential interactomics on a shared scale resolved the expansion into a coherent loss of transcription-activation machinery and a broad proteostatic and synaptic gain. Second, the calibrated posterior with the decision-theoretic call placed both arms on a common confidence scale: the 205 mHTT-gained contacts were the larger and the more robust arm (182 survive the stricter maximum-a-posteriori classification), and the 70 losses, though fewer, were comparably well-supported (45 of 70), identifying the expansion-gained proteostatic and synaptic interactions as genuine condition-dependent engagement rather than aggregation artefacts. Third, it improved cross-study reproducibility. The four source studies overlap only weakly at the naive-threshold level (mean pairwise Jaccard 0.19 across study pairs), so a conventional “reproducible in ≥ 2 studies” consensus recovers only ≈ 21% of the high-confidence interactors (*P* > 0.95). The remaining ≈ 79% are individually under-powered, credible in at most one study on its own; yet the per-study effect estimates show no cross-study heterogeneity (Cochran’s *Q* and the *I*^2^ statistic^1415^: 95% of calls homogeneous at *p* ≥ 0.05, median *I*^2^ = 0; Figure 3 C–D), so the integrated model recovers consistent, individually sub-threshold signal rather than study-specific noise — the expected gain from pooling under-powered studies. The per-study differential signal is individually too weak for any conventional test; because the datasets overlap little, the integrated model borrows strength across them to recover calls no single-study rule does. These analyses are a methods demonstration; the detailed biology is reported in the companion paper^7^.

**Figure 3.**
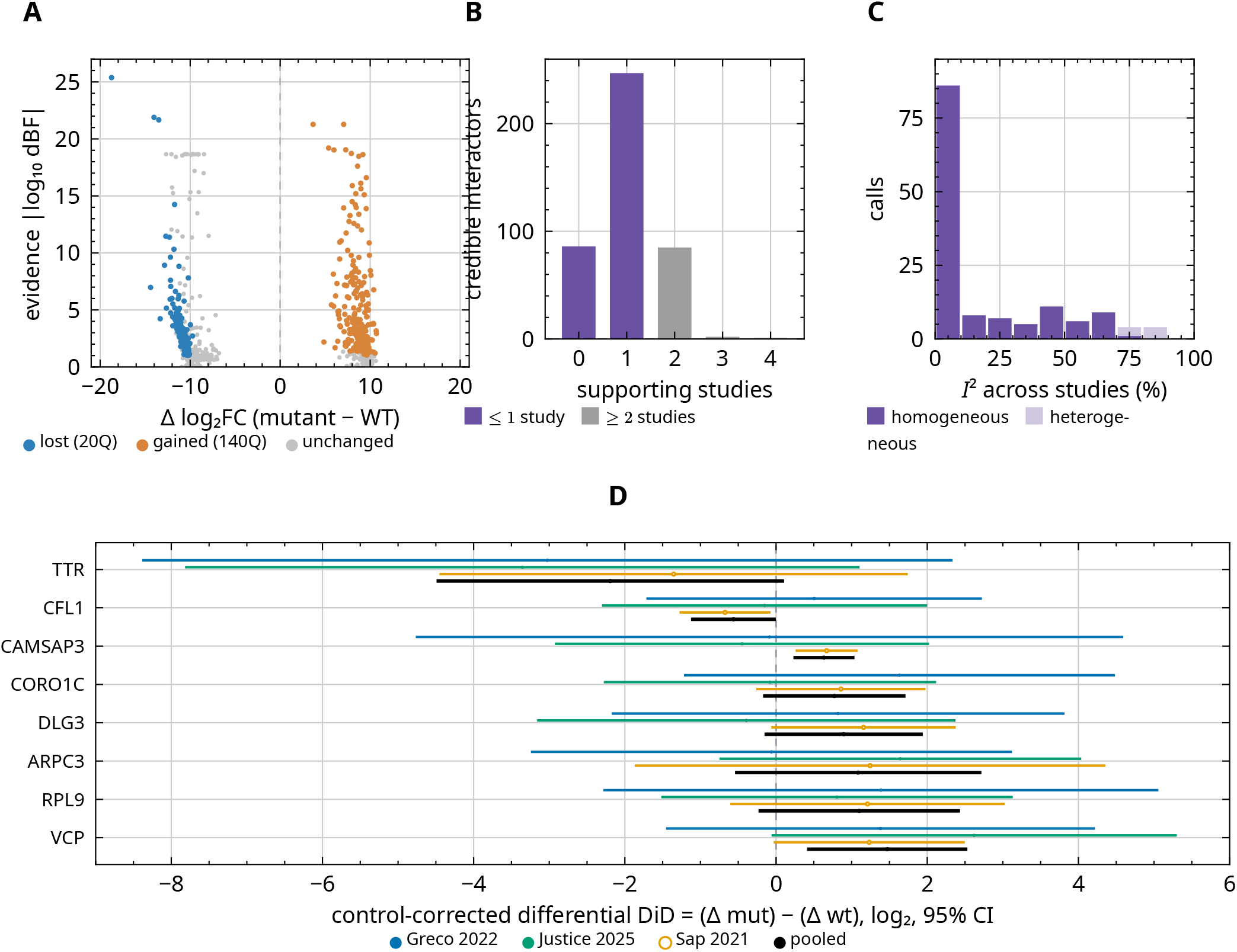
Three interactome analyses BayesInteractomics provides that no prior tool does. (A) Calibrated differential interactomics (Greco 2022 FLAG-HTT, 20Q versus 140Q at 10 months): signed differential evidence log_10_ dBF (positive = lost in the mutant; negative = gained) versus effect size |Δlog_2_FC|, with a calibrated differential posterior and Bayesian false discovery rate (BFDR) and the call gated on |log_10_ dBF| > 1 — none of SAINTexpress, CompPASS or MiST provides differential calibration. (B) The 421 credible interactors (P > 0.95) binned by supporting source studies under a conventional rule (detected in ≥ 50% of a study’s bait IPs AND ≥ 2-fold over controls): 79% rest on at most one study, so a naive “reproducible in ≥ 2 studies” consensus keeps only 88 (20.9%). (C, D) The calls are nonetheless cross-study homogeneous, not noise. Over the 140 calls testable in ≥ 2 co-detecting studies, the per-study control-corrected difference-in-differences (DiD) has median *I*^2^ = 0 and 95% show no significant heterogeneity (Cochran’s *Q*^1415^, *p* ≥ 0.05; 91% on raw values) (C); per-study CIs for representative calls are wide and cross zero (under-powered) yet overlap and agree in sign (D; black = inverse-variance pooled; Sap 2021 bait-only, hollow). A fair Stouffer^16^ / vote meta-analysis still recovers 0 of 275 calls at false discovery rate (FDR) < 0.05^17^ — single-study under-power, not contradiction; the integrated model borrows strength across studies. All panels are BayesInteractomics-internal (no SAINTexpress/CompPASS/MiST); B–D re-frame the companion meta-analysis (^7^, Suppl. Figs. 8–9). Source data `figure_differential_source_data.xlsx`, `figure_integration_recovery_source_data.xlsx`, `figure_homogeneity_source_data.xlsx`.

A second companion study applied BayesInteractomics to the obligate HTT–HAP40 complex, where its differential interactomics resolved stoichiometry-dependent interactor remodelling and the structural-docking step established that most candidate partners cannot engage the assembled complex because their interfaces are masked, separating direct binders from co-purifying partners^8^.

## Discussion

BayesInteractomics recasts AP-MS interactome calling as an end-to-end probabilistic inference: a data-driven prior — a deep neural network whose structural-contact prediction is combined with orthogonal sequence-, domain- and network-derived evidence by a calibrated meta-learner — conditions a multi-evidence Bayesian model fusing prey enrichment, bait–prey abundance correlation, and detection frequency through two complementary likelihoods reconciled by Bayesian model averaging (BMA), yielding a posterior controlled at a BFDR with native machinery for differential interactomics, per-call uncertainty, and an optional structural-docking refinement. To our knowledge no existing AP-MS scorer assembles a data-driven prior, multi-stream evidence fusion, calibration, uncertainty, differential testing, and reference-free structural arbitration into one coherent inferential object. This reframes how the score is used: not as a relative ranking alone, but as a per-call measure of confidence for prioritising experiments. On synthetic truth a posterior of *P* ≈ 0.8 recovers roughly four in five genuine interactors; on real data the posterior orders candidates by relative confidence and, once restricted to the detectability-adjusted positives the assay can support, matches the strongest comparator — so experimental validation can be prioritised by estimated confidence rather than truncated at an arbitrary rank.

We deliberately do not claim superior raw accuracy on a curated reference. Each of SAINTexpress, CompPASS and MiST is a capable single-condition scorer, and on a real human panel scored against CORUM ∪ IntAct BayesInteractomics is comparable to, not better than, all three; coverage-matched it ranks second behind MiST (Figure 2). This parity is expected, not a deficiency, because such a benchmark cannot adjudicate accuracy among AP-MS scorers for three measurable, structural reasons: it is open-world (a missing annotation reflects what has been tested, not what interacts, capping precision *a priori*^52^); it is partly circular (large tranches of curated edges were called by SAINT/CompPASS-class pipelines and deposited into the databases used as truth, tilting the reference toward the comparators^4^); and it has a hard recall ceiling (42% of reference partners carry no usable enrichment signal, forced false negatives for every method; Figure 2D^3^). A second, independent human dataset in a contrasting overexpression regime (Hein 2015; Extended Data Fig. 5) reproduces the pattern, so the parity reflects the benchmark, not the method. We therefore rest the reliability case not on this benchmark alone.

That reliability is established independently of curated catalogues. On synthetic data with complete, unbiased truth the raw posterior is discriminative but over-confident, and near-perfectly calibratable given ground-truth labels (ECE 0.15 → 0.015, ranking invariant) — a probabilistic structure no point-score comparator offers. On a fairly calibrated synthetic substrate BayesInteractomics is competitive with or ahead of all three comparators, first on AUROC in every batch with every paired confidence interval above zero, even with its prior switched off. Most consequentially, BayesInteractomics models the across-condition comparison directly: it contrasts conditions in the Bayes-factor domain, propagates per-condition uncertainty into a calibrated differential posterior, and ranks proteins by decision-theoretic validation priority — native differential interactomics on a shared probability scale that none of SAINTexpress, CompPASS or MiST provides (MiST is per-bait multi-evidence but, like the others, offers no across-condition contrast). In the companion HTT meta-analysis^7^ this raised cross-study reproducibility from ≈ 21% to 79% and recovered condition-dependent calls invisible to any per-study test, and it underpins the stoichiometry-dependent interactor remodelling resolved in the HAP40 study^8^. The docking step is a reference-free arbiter that sidesteps circularity and the recall ceiling, and method-agnostic cross-study reproducibility is the strongest real-data evidence.

These strengths carry honest limits. The method is species-general by construction — the DNN prior spans many organisms and the meta-learner featurises its transferred-STRING and Pfam channels per species, so a new organism needs only its own STRING and Pfam resources, not retraining — but what scales with annotation coverage is the prior’s *informativeness*: on sparsely annotated taxa it contributes little and calls are best read as evidence-driven, while the multi-evidence posterior remains intact. Absolute probability calibration on real data is not guaranteed: we therefore read the posterior and its BFDR as ranking and relative-confidence instruments rather than literal long-run frequencies, and report this as a limitation. Docking refinement is forward-looking and user-mediated. The real-data accuracy benchmark, for the reasons above, can corroborate but cannot rank, and the per-bait single-bait run model means multi-condition studies are assembled from independent runs. Within these bounds, the uncertainty-aware, differential core of BayesInteractomics delivers what point-score callers cannot — a reliable probabilistic ranking, explicit per-call uncertainty, and native across-condition inference — precisely in the regime where biological ground truth is genuinely unobtainable and reliable interactome calls matter most.

## Methods

### Machine learning

#### Data set generation & curation

Datasets of interacting and non-interacting protein pairs were generated by querying the RCSB Protein Data Bank (PDB). We included all available structures containing at least two distinct protein chains. For each structure, the Euclidean distance *d*_*i*_,_*j*_ between the *α*-carbon (*C*_*α*_) of every residue in one chain and all *C*_*α*_ atoms in the other chains was computed. A pair of protein chains was classified as **direct** interactors if the minimum calculated distance between them was below 5Å. Conversely, pairs within the same structure whose minimum *C*_*α*_ distance was above 5Å were classified as non-interacting. Protein sequence (Prot5) and cross-species network embeddings are retrieved from the STRING database^1819^.

With curated pairs defined, we next ensured non-redundant evaluation. The partitioning of the datasets into training (70%), validation (20%), and test sets (10%) was guided by minimising the mean cosine similarity of the protein embeddings between the sets — a similarity-minimised held-out construction that keeps highly similar pairs from leaking across folds. The selected DNN was trained on this single 70/20/10 partition. To bound the robustness of this single-split result, a 5-fold stratified cross-validation harness re-trains the same architecture on stratified resamplings of the curated set; across folds the test-set ECE standard deviation is 0.012 and the test-set ROC-AUC standard deviation is 0.007. A 5-model deep ensemble averaging the predictions of the five fold-checkpoints reaches test ROC-AUC 0.821 and ECE 0.044, confirming that the selected single-split model sits within the ensemble’s robustness band.

#### Training & Hyperparameter search of deep learning models

All deep-learning models were implemented in Julia using the Flux.jl library^20^. We trained a DNN on the concatenated protein embeddings to predict binary protein-protein interactions. The model was trained by minimising the binary focal loss function^21^ with the focusing parameter *γ* using either Adam or RmsProp optimiser.

The learning rate *η*(*t*_*e*_) was scheduled using cosine annealing^22^ with a linear warmup period. The schedule for epoch *t*_*e*_ is defined by:

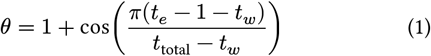

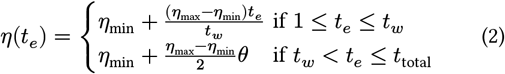

with the maximal learning rate *η*_max_, the minimal learning rate *η*_min_, the total number of epochs *t*_total_, and the number of warmup epochs *t*_*w*_.

To regularise the DNN and smooth the embedding space, Mixup^23^ was applied to generate virtual training samples. Given two randomly selected training samples (*x*_*i*_, *y*_*i*_) and (*x*_*j*_, *y*_*j*_), a virtual samples 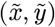 is generated through convex combination:

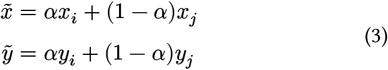

As an additional data augmentation step, each training pair (*A, B*) was randomly swapped to (*B, A*) with a 50% probability to enforce permutation invariance.

The optimal model configuration was identified through three successive stages of random search, with each stage using a progressively narrower search space centred on the best-performing configurations from the previous stage (Table 2). In total, 1100 configurations were evaluated. The performance of each configuration was assessed on the validation set using the MCC as the primary selection metric.

**Table 2:**
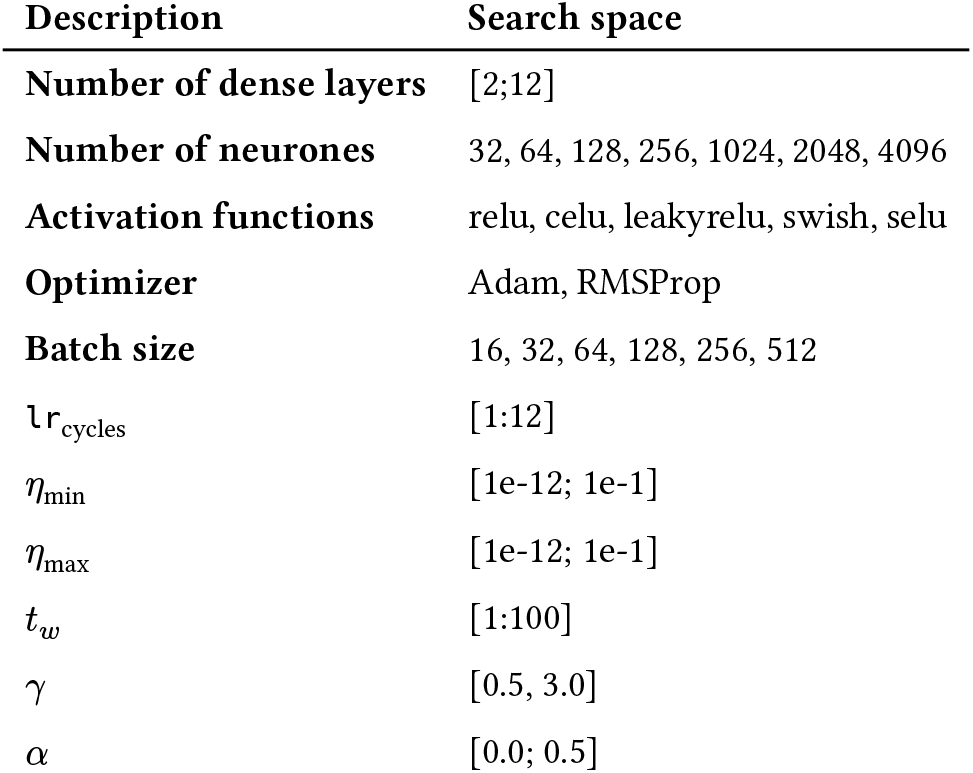
Hyperparameter search ranges.

The configuration selected by this search defines the trained DNN the meta-learner stack described below consumes its output as a fixed deep-learning prior column, so all downstream meta-learner performance is reported conditional on this single trained network.

#### Meta-learner

To integrate the DNN prior with additional sources of evidence, the meta-learner is implemented as a stacking ensemble^24^ of six fixed-hyperparameter base learners — histogram gradient boosting, EvoTrees, L2-regularised logistic regression, *k*-nearest neighbours, random forests, and extra trees — whose out-of-fold predictions are blended by an L2-regularised logistic-regression layer. A post-hoc isotonic calibrator is then fitted on a held-out validation slice that the stack itself has not seen, so calibration metrics reported on the test set are unbiased.

The stack consumes 14 raw input features grouped into four families: the seven STRING evidence channels in-species^1819^ (neighborhood, fusion, phylogenetic, coexpression, experimental, database, textmining), four cross-species transferred STRING channels obtained by mapping the same evidence types across orthologous proteins, two Pfam protein-domain-interaction features summarising whether the two proteins’ Pfam domain pair has any known interaction and how many, and the DNN prior probability. The transferred-STRING and Pfam domain features are computed per species from each organism’s own STRING and Pfam resources, so the meta-learner is not tied to a single reference proteome and scores a non-human bait on species-correct evidence rather than on human-derived or zero-filled features. We report two variants of this schema: a network-only variant using the 14 raw inputs directly, and an uncertainty-augmented variant that adds a fifteenth column — the Monte-Carlo-Dropout standard deviation obtained from *K* = 30 stochastic forward passes through the DNN — exposing the network’s per-pair predictive uncertainty to the blender. The network-only variant is the production default; the uncertainty-augmented variant is retained as an alternative whose relative performance is reported in the Results.

Tuning uses Brier loss as the scoring measure under 5-fold group-disjoint cross-validation inside the MLJ.jl framework. Both variants achieve a calibrated test-set ECE at or below the 0.035 acceptance threshold we adopted for downstream use.

Examining the blender’s coefficient magnitudes on standardised inputs (Extended Data Fig. 1), the seven in-species STRING channels collectively account for only ∼ 20% of the blender’s direct importance, while the DNN prior together with the four base-model out-of-fold predictions account for ∼ 78% (panel B), the small remainder carried by the transferred-STRING and domain features. This ∼ 20% figure is an upper bound on the direct contribution of STRING evidence: each of the base learners internally also consumes the same in-species STRING channels, so a portion of the base-prediction family’s importance traces back to STRING processed non-linearly inside the base models. The headline conclusion is robust regardless of accounting choice — direct STRING evidence is a minor contributor at the blender layer, with the bulk of the discriminative signal flowing through the DNN prior and the stacked base learners; the four transferred-STRING channels in particular carry negligible direct weight even when computed species-correctly, confirming that cross-species transfer adds little once the in-species evidence, the base learners and the deep-learning prior are present.

A feature-family ablation (Extended Data Fig. 2) shows that the selected transferred-STRING-plus-domain schema attains the highest held-out ROC-AUC of the eight feature configurations evaluated, whereas adding a subcellular-localisation family on top of it lowers ROC-AUC and raises calibrated ECE. The selected configuration therefore sits at the discrimination elbow, beyond which further feature families yield no AUC gain; Gene-Ontology semantic-similarity features, evaluated separately, were likewise not retained in the production schema.

### Bayesian Inference

To analyse the experimental evidence, three different models were implemented:

1. a three-level hierarchical Bayesian model to robustly quantify protein enrichment
2. a hierarchical Bayesian linear regression to assess the correlation between prey and bait protein abundances,
3. a Bayesian Beta-Bernoulli model to assess whether a protein was detected more frequently in sample versus control replicates.

The evidence from these three models was then combined using the two-component mixture model described below.

#### Hierarchical Enrichment Model

To quantify the enrichment of prey proteins, we developed a three-level hierarchical Bayesian model that shares information across different experiments and protocols, yielding more robust estimates. The abundance of each protein in the sample and control groups was assumed to be normally distributed.

The model estimates a separate mean abundance (*μ*) and precision (*ρ* = 1/*σ*^2^) for each individual experiment. These experiment-level means are modelled as being drawn from a protocol-level distribution, which is in turn drawn from a global, top-level distribution. This hierarchical structure enables partial pooling of information, allowing experiments with less data to borrow strength from the overall dataset.

Weakly informative, empirical priors were used for all top-level parameters. The hyperparameters for the global mean prior, 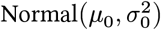, were estimated from the data: *μ*_0_ was set to the global mean intensity across all proteins and samples, and 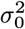 was set to the global mean variance. Similarly, the hyperparameters for the Gamma prior on the precision terms were estimated from the variance of all measurements. This data-driven approach anchors the priors in the observed distribution of the data while still allowing for robust Bayesian updating. The likelihood of the observed protein abundances was modelled as a Normal distribution with the corresponding experiment-level mean and precision.

Explicitly, the observed log2-intensities follow 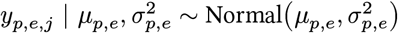, with the experiment-level means drawn from *side-specific* protocol-level distributions for the control and sample arms, 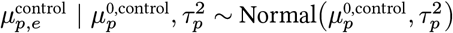 and 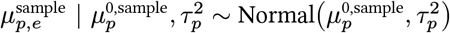 so that the two arms are fitted as separate protocol-level means rather than as a single mean with an added effect term. The protocol-level baselines themselves draw from a global Normal whose hyperparameters *μ*_0_ and 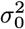 are computed directly from the data (the grand mean and grand variance of all log2-intensities), while the within-protocol experiment precision *τ*_*p*_ and the per-experiment observation precision 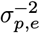 carry Gamma priors on the precision whose shape and scale parameters are themselves fit by maximum likelihood to the empirical precision distribution of the input matrix — a data-anchored, weakly-informative scheme that lets the data dominate the posterior without privileging a fixed variance prior, with a Gamma(1, 1) fallback when the empirical fit fails. The protocol-level log2-fold-change is not a latent parameter of the model but is computed post-hoc as the posterior-mean difference 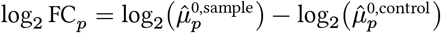, and the enrichment Bayes factor is the posterior-odds-to-prior-odds ratio for the one-sided hypothesis *H*_1_ : log_2_ FC_*p*_ > 0 against *H*_0_ : log_2_ FC_*p*_ ≤ 0 evaluated on this transformed quantity. Posterior inference is performed by variational message passing in RxInfer.jl^25^, the same engine used for the regression sub-model.

From this hierarchical posterior we derive a per-condition enrichment Bayes Factor, BF_enrichment_, comparing the hypothesis of meaningful enrichment of a prey protein in the sample relative to the control against the null. We interpret all Bayes factors reported in this work on the Kass-Raftery evidence scale^26^, with BF ≤ 3 corresponding to “barely worth more than a bare mention”, 3 < BF ≤ 20 to “positive”, and higher tiers to progressively stronger support; this same scale frames the threshold-based empirical-null assignment used in the two-component mixture model below.

#### Hierarchical Regression model

To model the relationship between the abundance of a prey protein and the bait protein, we implemented a hierarchical Bayesian linear regression model with RxInfer.jl^25^. This approach allows for the estimation of a separate regression for each experimental protocol while sharing information between them, providing more robust estimates.

For each protocol *j*, the relationship was modelled as *y*_*i*_ = *β*_*j*_ + *α*_*j*_*x* + *ε*_*j*_, where *y*_*i*_ is the abundance of the prey protein, *x* is the abundance of the bait protein, and *ε*_*i*_ represents the residual error. The protocol-specific intercepts (*β*_*j*_) and slopes (*α*_*j*_) were modelled as being drawn from common global distributions:

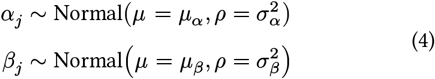

Weakly informative hyperpriors were placed on the global mean and precision parameters (*μ*_*α*_, *μ*_*β*_, *ρ*_*α*_, *ρ*_*β*_). Specifically, the prior for the global analysis slope, *μ*_*α*_, was a Normal(*μ* = 0, *ρ* = 0.153) distribution, chosen to place 95% of the prior mass on slopes between −0.3 and 0.3. The prior on the global intercept *β* was set to the global mean intensity across all proteins and samples, and 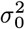 was set to the global mean variance. The Gamma distribution Γ(*α* = 5, *θ* = 2) with shape parameter *α* and scale parameter *θ* was assigned as the prior for the residual precision of the model.

Two observation models are available for the residual term: a Normal likelihood, 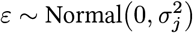, and a robust Student-*t* likelihood, 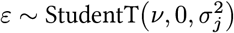 (the default), so that outlying replicates do not unduly dominate the slope estimate. When model comparison is enabled, BayesInteractomics fits both observation models and selects the one minimising the widely applicable information criterion (WAIC)^2728^,

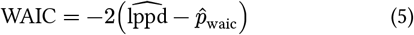

where 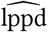 is the estimated log pointwise predictive density and 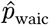 the effective-parameter penalty; the lower-WAIC variant is adopted automatically. When the Student-*t* model is selected, its degrees of freedom *ν* are themselves optimised over the interval [5, 50] by Brent’s-method minimisation of the WAIC. The slope carries, by default, a Jeffreys-Zellner-Siow (JZS) mixture-of-*g*-priors^2930^ expressed in the Normal-Gamma scale-mixture form,

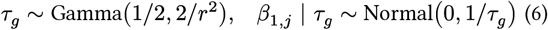

with scale 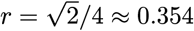 (the JASP convention); marginalising *τ*_*g*_ recovers a Cauchy(0, *r*) slope prior, but the scale-mixture parameterisation is retained because it admits tractable variational message passing in RxInfer.jl^25^ whereas the marginalised Cauchy does not, and setting the scale to zero falls back to a flat Normal slope prior for backward compatibility. The per-protein residual precision is estimated empirically when a protein has at least five observations and otherwise shrunk towards the protocol-level pooled precision; a posterior-variance floor of 0.01 guards against variational over-confidence in sparsely observed cells.

The Bayes Factor BF_regression_ was computed to test the one-sided hypothesis that the slope is meaningfully positive, *H*_1_ : *β*_1_ > 0.1, against the null hypothesis *H*_0_ : *β*_1_ ≤ 0.1. The value 0.1 here is a calibration-anchored decision boundary operationalising a biologically meaningful positive slope — not a prior parameter (in particular it is distinct from the *α* shape of the residual-precision Gamma prior above).

#### Bayesian Beta-Bernoulli Model

To assess the evidence for specific detection of a prey protein in the presence of the bait, we employed a Bayesian Beta-Bernoulli model. The detection of a prey protein in any given replicate was modelled as a Bernoulli trial, where the probability of detection in the sample and control groups is denoted by *θ*_*s*_ and *θ*_*c*_, respectively.

We placed a weakly informative Beta distribution, Beta(*α* = 3.0, *β* = 3.0), as a priror on both *θ*_*s*_ and *θ*_*c*_. Given the conjugate nature of the Beta-Bernoulli model, the posterior distribution for each probability was calculated analytically. For the sample group, the posterior is

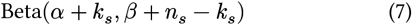

where *k*_*s*_ is the number of replicates where the prey was detected in the sample group and *n*_*s*_ is the total number of sample replicates. The posterior for the control group was calculated analogously:

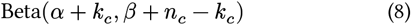

where *k*_*c*_ and *n*_*c*_ are the detection counts and total number of replicates for the control group, respectively. The Bayes Factor (BF_Detection_) was computed by comparing the posterior distributions via simulation to estimate the probability that the detection rate in the sample group is greater than in the control group.

#### Copula evidence-density mixture

To combine the three independent streams of experimental evidence (enrichment, correlation, and detection), we model their joint per-protein density on the log-Bayes-factor scale (*b*_*e*_, *b*_*c*_, *b*_*d*_) = (log BF_enrichment_, log BF_regression_, log BF_Detection_) as a three-component mixture. The three components represent a background null (*H*_0_, non-interactors), an anchored agnostic class capturing proteins with ambiguous evidence, and an interactor alternative (*H*_1_, true direct interactors), so that 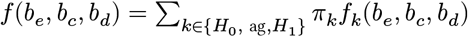. Within each component the joint density is constructed via Sklar’s theorem^31^ as a single three-dimensional copula coupling three univariate evidence marginals (a single copula per component, not a vine):

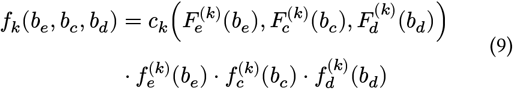

where *c*_k_ is the component copula density and 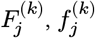 are the marginal cumulative and density functions for evidence dimension *j* ∈ {*e, c, d*}.

For the *H*_0_ and *H*_1_ components the copula *c*_*k*_ is selected by minimising the Bayesian information criterion (BIC)32, 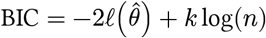 with a single shape parameter (*k* = 1) per family, over exactly four single-parameter families — Clayton, Frank, Gumbel, and Gaussian — covering lower-tail, no-tail symmetric, upper-tail, and weak-symmetric dependence regimes respectively. The anchored agnostic class uses a fixed independence (Gaussian, identity-correlation) copula with Normal marginals whose enrichment mean is anchored at zero to prevent redundancy with the *H*_0_ class. The *H*_1_ enrichment marginal is a location-shifted Gamma anchored at the Jeffreys evidence threshold 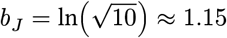 nats, giving zero *H*_1_ density below this threshold and a smooth sigmoid transition above it; the remaining marginals are Normal with a Kolmogorov–Smirnov-checked auto-upgrade to a Student-*t* location-scale form. Family selection is not a one-shot decision: after a five-iteration burn-in the *H*_0_ and *H*_1_ copulas are re-fit and re-selected as the expectation-maximisation (EM) responsibilities reallocate proteins between classes. If fewer than fifty observations are available for a component, or if all four families fail to fit, that component falls back to the independence copula.

The mixing weights 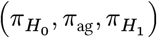 carry a Dirichlet prior whose mass on 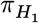 is literature-anchored to empirical AP-MS hit-rate distributions, with a default Beta(80, 20) marginal on 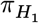 reflecting the prior belief that true direct interactors constitute a minority of all identified proteins^433634^ (overridable per experiment type). This prior is not co-tuned with the DNN prior or any other component of the model, so component-ablation studies that drop the DNN prior remain uncontaminated. Fitting uses a MAP-EM algorithm with twenty multi-restart initialisations (quantile-shift, *k*-means, and random-perturbation strategies) and SQUAREM acceleration35 with step-halving guards. The per-protein copula sub-model Bayes factor BF_copula_ is the posterior-odds-to-prior-odds ratio for the interactor class, which is subsequently averaged against the latent-class sub-model below.

#### Latent-class factored-marginal mixture

As a dependence-free counterpart to the copula sub-model, we fit a three-component finite mixture^36^ on the same joint per-protein log-Bayes-factor triplet, sharing the identical *H*_0_ background / anchored agnostic / *H*_1_ interactor decomposition but replacing the copula coupling with a conditional-independence assumption: within each class *k* the three evidence dimensions are conditionally independent and the joint density factorises as 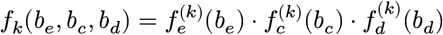. This factorisation makes the likelihood a tractable product of one-dimensional densities and removes any dependence-misfit risk at the cost of ignoring genuine within-class rank correlation — the opposite trade-off to the copula sub-model.

The *H*_0_ class uses a heavy-tailed Student-*t* enrichment marginal with degrees of freedom *ν* ∈ {3, 5, 7, 10} selected by BIC, giving robustness to outlier quantification artefacts; the anchored agnostic class uses a Normal marginal whose mean is fixed at zero to prevent redundancy with the null; and the *H*_1_ interactor class uses a right-shifted enrichment marginal drawn from one of {Gamma, LogNormal, Weibull} — allocated across restart thirds and confirmed by BIC at iteration five — gated through a sigmoid transition *σ*(*w*(*b* − *b*_*J*_)) at the Jeffreys evidence threshold 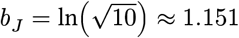 nats^37^. Detection marginals are nonparametric (discrete empirical distributions) in all three classes and correlation marginals are Normal with a Kolmogorov–Smirnov-checked auto-upgrade to a Student-*t* location-scale form when uniformity of the probability-integral transform fails.

The mixing weights 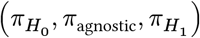 carry a Dirichlet prior whose concentration *α* is estimated by Empirical Bayes (EB) via the Minka fixed-point iteration^38^, with two-stage post-convergence clamping (total concentration rescaled to [3, 30], individual components floored at 0.5); this is distinct from the copula sub-model’s Beta(80, 20) weight prior. Inference is MAP-EM^39^ with convergence at relative log-likelihood change below 10^−6^ or two hundred iterations, a step-halving monotonicity guard that activates after iteration ten (the copula sub-model’s SQUAREM acceleration is not used here), twenty multi-restart initialisations, and optional one-percent-tail winsorisation. A small number of post-EM refinements — a monotonicity floor on log-likelihood ratios, deterministic annealing on bimodal detection evidence, marginalisation over a nine-point EB prior grid, and a Kullback– Leibler merge of the *H*_0_ and agnostic classes when they collapse onto the same distribution — stabilise degenerate fits and are detailed in the Extended Data. Per-protein soft responsibilities *γ*_*n,k*_ are exported alongside the sub-model Bayes factor

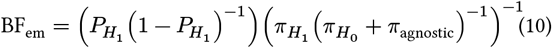

— the per-protein posterior odds of the interactor class divided by the prior odds derived from the EM-estimated mixing weights — which is then averaged with the copula sub-model’s Bayes factor in the Bayesian Model Averaging step.

Both sub-models are fit by MAP-EM39 and share a common set of numerical safeguards, differing only in how they stabilise the iteration. Convergence is declared once the relative change in log-likelihood between successive iterations drops below 10^−6^, or after a ceiling of two hundred iterations per restart; a restart that reaches the ceiling without satisfying the tolerance is flagged non-converged, and the best restart is selected by maximum converged log-likelihood across twenty multi-restart initialisations spanning three strategy families (quantile-based, *k*-means, and random Dirichlet-perturbation seeds), which together protect against local optima. The two sub-models then diverge in their acceleration: the copula sub-model additionally applies SQUAREM acceleration35 on the best restart, whereas the latent-class sub-model relies instead on the step-halving monotonicity guard described above — activating after iteration ten and reverting any M-step that decreases the log-likelihood — and does not use SQUAREM. The two sub-models also carry distinct mixing-weight priors: the Beta(80, 20) marginal regularises the copula sub-model’s interactor weight 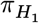 (Section above), while the latent-class sub-model carries a separate Empirical-Bayes Dirichlet on its 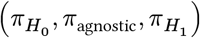 weights whose concentration is estimated from the data by the Minka fixed-point iteration38 and clamped post-convergence.

#### Bayesian Model Averaging

Rather than committing to either sub-model, BayesInteractomics produces its final per-protein interaction call by BMA40 over the two three-component views of the joint per-protein log-Bayes-factor triplet: the dependence-aware copula evidence-density mixture (which couples the enrichment, correlation, and detection marginals through a single three-dimensional copula per class) and the factored-marginal latent-class mixture (which assumes those three dimensions are conditionally independent within each class). Both sub-models share the identical *H*_0_ / anchored-agnostic / *H*_1_ decomposition — they differ only in how they model within-class dependence, not in their component count — so averaging them hedges the dependence-aware view (which can overfit when within-class rank correlation is noisy) against the dependence-free view (which can under-fit when there is genuine cross-evidence corroboration), with the two views genuinely disagreeing in the middle-evidence band.

The sub-models are combined by leave-one-out (LOO) stacking^41^. For each protein *i* and sub-model *k*, the LOO predictive density 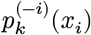 is estimated by Pareto-smoothed importance sampling (PSIS)^28^, and the stacking weights are obtained as

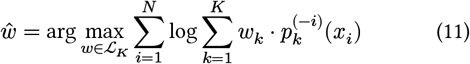

subject to *w*_*k*_ ≥ 0.05 for every sub-model — a five-percent weight floor that prevents winner-take-all degeneracy when one sub-model fits marginally better but both remain reasonable. The per-protein PSIS reliability statistic, the Pareto 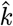 diagnostic, is reported for every call 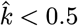 reliable, 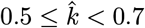 borderline,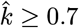 unreliable).

The combined Bayes factor is the stacking-weighted **linear** average of the sub-model Bayes factors,

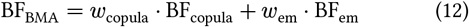

Pooling the Bayes factors directly, rather than back-deriving the combined factor from an averaged posterior, keeps the result insensitive to the choice of prior odds. The combined posterior probability is then recovered from the pooled Bayes factor and the shared EB prior odds,

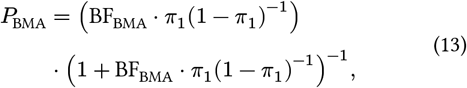

where *π*_1_ is the EB posterior-mean weight of the interactor class. To keep downstream interpretation transparent about which sub-model is driving each call, the analysis output exposes the raw sub-model Bayes factors, the per-sub-model stacking weights, the reliability statistic, and a per-protein model-disagreement indicator. The disagreement flag is set when the two sub-models reach **opposite** binary classifications at the *P* > 0.5 decision boundary — that is, when (*P*_copula_ > 0.5) ≠ (*P*_em_ > 0.5) — flagging proteins where the two views qualitatively disagree on whether the protein is an interactor; it is not a function of the magnitude of the Bayes-factor difference between the sub-models.

This averaging step is distinct from, and sits one level above, the **internal** copula-mixture combining of the three evidence streams. That internal operation lives inside the copula sub-model, where the enrichment, correlation, and detection evidence are fused into one composite at the joint-density level via a single three-dimensional copula; BMA then averages the resulting copula-combined output against the latent-class-combined output. The two operations are not the same: one combines evidence dimensions within a sub-model, the other averages whole sub-models.

#### Decision thresholds and FDR control

The operational decision tiers reported throughout this work — a Bayesian FDR of BFDR ≤ 0.05 (“significant”) and BFDR ≤ 0.01 (“strong”) — are derived from the synthetic calibration sweep rather than chosen ad hoc. The same 5 × 5 grid over interaction prevalence 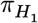 and effect-size scale used to calibrate the BMA posteriors (ten replicates per grid point and *N* = 10^4^ synthetic proteins per replicate, for two hundred and fifty scenarios in total) carries known ground-truth labels, so for any candidate posterior cut-off *t* both the declared BFDR and the empirical false-discovery rate can be evaluated against truth in each scenario:

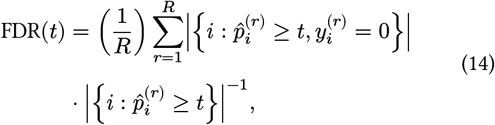

averaging over the *R* replicates of a scenario. The reported thresholds correspond to the operating points at which the empirical FDR’s 97.5th percentile across all two hundred and fifty scenarios remains within the declared BFDR band4243, so that the nominal 0.05 and 0.01 tiers are conservative with respect to the simulated truth rather than nominal-only. The full empirical false-discovery and declared-BFDR curves (one hundred grid points each) are exposed in the analysis output, so users may select alternative operating points matched to their own error tolerance.

#### Output probability calibration

Before the operational decision tiers above are applied, the per-protein posterior probabilities emitted by the BMA step are recalibrated against synthetic ground truth. This downstream recalibration is distinct from the isotonic calibration applied inside the meta-learner’s stacking ensemble (Section “Meta-learner”): that earlier step calibrates the meta-learner’s stacked class-probability output — the prior — at the feature level, whereas the step described here recalibrates the final combined Bayesian posterior — and its derived BFDR — at the per-protein decision level. Calibration follows a three-step protocol. First, the procedure regenerates the same synthetic backbone used to set the decision thresholds: a 5 × 5 sweep over interaction prevalence 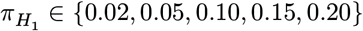 and effect-size scale {0.5, 0.75, 1.0, 1.25, 1.5}, with ten replicates per grid point and *N* = 10^4^ synthetic proteins per replicate, for two hundred and fifty scenarios in total, drawing Bayes-factor triplets from the fitted latent-class distributions with the corresponding ground-truth labels. Second, a Platt-scaling map^44^ is fit in logit space,

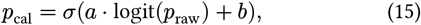

with parameters (*a, b*) optimised by minimising binary cross-entropy against the synthetic labels. Third, a ten-fold stratified cross-validated ECE^45^,

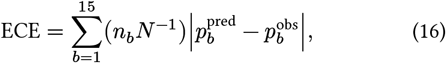

evaluated over fifteen equal-width probability bins (with *n*_*b*_ the count in bin 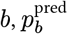 the mean predicted probability, and 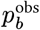 the empirical positive fraction), acts as a safety guard: the transform is adopted only if the mean cross-validated ECE across the ten folds falls below 0.05, and otherwise the raw posteriors are retained and a warning badge is surfaced in the diagnostic report. Bayesian false-discovery rates^46^ are then computed from the calibrated (or, when the guard rejects the transform, the raw) posteriors,

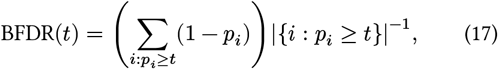

and it is from this calibrated FDR / sensitivity curve that the BFDR ≤ 0.05 and BFDR ≤ 0.01 operating tiers of the preceding subsection are read off. Because this recalibration is fit on Bayes-factor triplets resampled from the model’s own fitted latent-class distributions rather than on external truth, it is a within-model recalibration validated in-distribution; it is not guaranteed to transfer to real data, whose dominant error modes — incomplete annotation, condition-specific background and control carry-over — the generator does not reproduce. We therefore treat the recalibrated posterior and its derived BFDR as ranking and relative-confidence instruments and report absolute probability calibration on real data as a stated limitation rather than a validated property (Results).

#### Sensitivity and diagnostic outputs

Three families of post-fit diagnostics quantify the reliability of each analysis run. *Posterior predictive checks* (PPC)47 re-fit the hierarchical enrichment and regression sub-models on the analysed proteins — by default all of them, since the per-run cap on the number of proteins checked acts as an opt-in rather than a default sample size — draw one thousand posterior-predictive samples per protein, and report a per-protein posterior predictive check (PPC) *p*-value as the proportion of simulated discrepancies more extreme than the observed one; a uniform aggregate distribution of these *p*-values indicates a well-specified model. *Residual diagnostics* compute standardised regression residuals 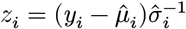, flag outliers at |*z*| > 3, and inspect normality via quantile-quantile plots and heteroscedasticity via scale-location plots. *Calibration assessment* re-uses the synthetic-anchored Platt model of the preceding subsection to produce a reliability diagram for each downstream classification strategy.

Two automatic *quality gates* check whether the mixture fit is well-posed before its posteriors are trusted. A Kolmogorov-Smirnov (KS) statistic per marginal flags goodness-of-fit failure (default warn at KS > 0.10, fail at KS > 0.15, the latter triggering automatic Student-*t* remediation of the offending component), and a per-stream Kullback-Leibler (KL) divergence flags insufficient component separation as a single binary verdict — each evidence stream passes when its KL divergence stays below 0.5, and component-level contamination passes only when all three streams pass, with failure indicating that two components have collapsed onto one another on at least one evidence dimension. The overlap coefficient (OVL) and within-class Spearman correlation are reported per component as descriptive separation summaries without warn/fail thresholds.

*Prior sensitivity* is quantified in four layers. First, the Dirichlet concentration *α* governing the three-component mixing prior is estimated by EB via the Minka fixed-point iteration^38^,

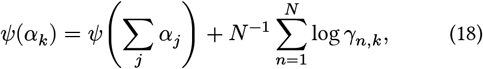

with two-stage post-convergence clamping: the total concentration ∑_*j*_ *α*_*j*_ is first rescaled to lie in [3, 30], and then each individual component is floored at 0.5, the two stages together keeping the prior weakly informative without letting any component collapse. Second, a nine-point constant-strength simplex grid is built around the EB *α* estimate and the per-grid-point posteriors are averaged with softmax-of-BIC weights,

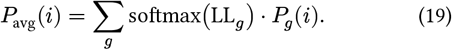

Third, a Cartesian product over the latent-class Dirichlet, copula EM Beta, and optional Beta-Bernoulli detection priors (roughly twenty-seven grid points) probes the joint sensitivity. Fourth, each protein receives a *classification stability* traffic light (robust / sensitive / fragile) indicating whether its *P* > 0.5 classification flips across the joint grid. This stability label, together with low-data flagging, residual-outlier status, PPC failures, and the BMA Pareto 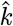 reliability statistic^28^, is aggregated into a single per-protein diagnostic flag (ok / warning / fail) emitted alongside every interaction call.

### Data preparation

#### Data curation

Raw protein-identifier columns are standardised in four sequential pre-analysis steps before any modelling. First, rows tagged as contaminants (“CON—” prefix) or decoy reverse hits (“REV—” prefix) are removed. Second, semi-colon-delimited protein-group identifiers are split into individual rows, each daughter row retaining the original quantification values, so that group-level identifications are propagated downstream as ambiguous candidates rather than discarded. Third, each identifier is resolved to its canonical UniProt accession and STRING preferred name via the STRING REST API^18^, with a per-species local cache — written beside the input data file as a per-species, per-data-hash artefact — preventing repeated network lookups on subsequent runs; identifiers that fail to resolve are recorded and retained unchanged. Fourth, identifier collisions — multiple input rows mapping to the same canonical accession — are merged into a single row by element-wise maximum (default) or arithmetic mean on the linear scale, optionally with interactive confirmation or with auto-approval that fires when all merge candidates in a group share the same leading *N*-character prefix. When a bait protein is named, its identity is tracked through all four steps and its post-curation row index is returned for downstream addressing. Every curation action is logged into a serialisable curation report whose replay mode reproduces an identical curation on new input without re-querying STRING — a property we exploit to guarantee bit-identical analyses across continuous-integration runs.

#### Input data quality control

Five automated quality control (QC) checks run on the curated, normalised log_2_-intensity matrix before any modelling, with an overall verdict reduced to the worst sub-flag under the precedence fail > warning > ok. The *scale* check flags suspected linear-scale data accidentally supplied in place of log_2_ values, emitting a warning when the maximum intensity exceeds 1000. The *replicate correlation* check computes pairwise Spearman correlations within each replicate group (per protocol and condition) over the proteins detected in both replicates of a pair, and flags any minimum pairwise correlation below 0.80 (warning) or below 0.60 (fail), the latter marking the empirical floor below which downstream Bayes-factor inference becomes unreliable. The *missingness asymmetry* check identifies replicates whose per-replicate missing-data fraction exceeds two times (warning) or three times (fail) the group-median missingness, catching replicates lost to acquisition issues, instrument drift, or bad samples. The *distribution shape* check inspects, per replicate, the bimodality of the non-missing intensities via low excess kurtosis (warning at < −1.2), a heavy right tail via high excess kurtosis (warning at > 7.0), and intensity-floor spiking — the fraction of values equal to the minimum non-zero value or to exactly zero — emitting a warning at > 20% and a fail at > 40% of values; skewness is computed and exposed alongside these statistics but does not itself contribute a flag. Finally, *PCA separation* scores Fisher’s discriminant ratio^48^ on the first two principal components,

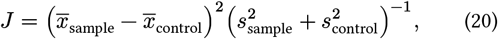

evaluated across cascading complete-case subsets (proteins present in 100%, then 80%, then 50% of replicates, taking the first threshold that yields at least twenty proteins; principal component analysis (PCA) is skipped entirely if fewer than twenty proteins pass at 50%). It emits a warning when both the PC1 and PC2 ratios fall below 1.0, or when PC1 explains less than 25% of the variance; PCA separation never escalates to a fail, and for multi-protocol datasets a strong per-protocol decomposition caps a weak combined PCA at warning to avoid over-flagging. Each check exposes its metric values and per-protocol decomposition in the analysis output, so that any warning or fail is actionable: the user learns which replicate, which check, and which threshold was crossed.

#### Input data normalisation

AP-MS intensity matrices are normalised on the log_2_ scale before any imputation, model fitting, or differential analysis is performed. Four strategies are available: no normalisation (for data already normalised upstream); row-centering (subtraction of the per-protein row mean across all samples, removing per-protein scale differences); median-of-ratios (DESeq-style size-factor normalisation^49^, with per-sample size factor

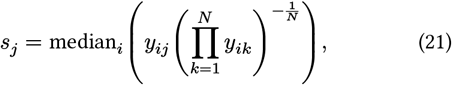

the geometric mean taken over the *N* samples in which protein *i* is fully observed); and a combined strategy (median-of-ratios followed by row-centering). By default, an automatic mode inspects the per-protocol intensity distributions and selects the combined strategy when a mismatch is detected, falling back to no normalisation otherwise. When a bait protein is named, an additional bait-anchoring step runs after normalisation and before the regression model: for each protocol *c* it subtracts from every prey the per-protocol offset

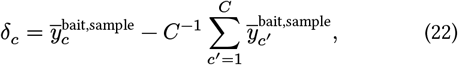

the difference between protocol *c*’s mean bait sample level and the grand mean of per-protocol bait sample means, so that protocol-level bait-abundance offsets do not inflate per-condition enrichment differences in differential interactomics. Normalisation, imputation, bait-anchoring and regression are enforced in that fixed order; calling imputation before normalisation would invalidate the per-column missing not at random (MNAR) dropout-curve fit described in the next subsection, whose intercept and slope assume an already-normalised intensity scale.

### Handling of missing data

AP-MS intensity matrices contain a mixture of two missingness regimes that we treat differently. Some non-detections are missing at random (MAR) — independent of the underlying abundance — while the majority follow a detection-limit pattern in which low-intensity peptides are systematically less likely to be detected (MNAR)^50^. Naive MAR-style imputation on MNAR cells imputes values that are too high and too confident, which propagates into downstream Bayes factors and produces saturation pathologies on real data.

We therefore use detection-limit-calibrated MNAR-aware imputation. For each mass spectrometry (MS) run column *c* we fit a logistic dropout curve to the observed detection pattern,

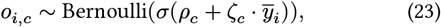

where *o*_*i,c*_ is the detection indicator for protein *i* in column *c* and 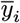 is the mean of log_2_-intensities for that protein across columns where it was detected. Each missing entry is then imputed by a single draw from the tilted Gaussian

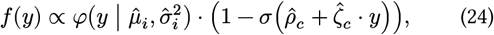

— a column-shared Normal centred at the per-protein mean 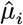 multiplied by the column’s non-detection probability^51^. The per-column empirical variance of the resulting post-imputation matrix, 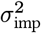, is injected into the Bayesian regression observation factor as an additive variance term on cells flagged as imputed, so the regression posterior is aware of which cells were imputed and how confidently. This single-draw scheme is the default; an optional multi-imputation mode with Rubin-style within-and-between pooling is available for sensitivity analyses.

### Differential and structural analysis

#### Differential interactome analysis

Because each BayesInteractomics run scores a single bait against its own control pool, condition comparisons are performed as a downstream contrast over two or more completed per-bait analyses rather than as a joint fit. Given two analysis results *A* and *B* for the same bait under different conditions (for example wild-type versus mutant, or two cell lines), prey proteins are matched by identifier across the detected sets, and the per-protein evidence is contrasted directly in the Bayes-factor domain. The differential Bayes factor (dBF) for a shared prey is the ratio of its per-condition copula Bayes factors, computed as a subtraction in log-space,

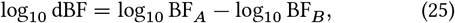

so that a positive value indicates stronger interaction evidence in condition *A*. The same ratio is formed separately for each evidence channel — enrichment, replicate correlation, and detection consistency — yielding per-channel differential Bayes factors that diagnose which line of evidence drives a given change. To keep the contrast on the interaction signal rather than on a condition-level prior, the per-condition posterior probabilities entering the comparison are re-derived from the copula Bayes factor alone (*p* = BF/(1 + BF)), discarding the meta-learner prior, which is shared across conditions for the same protein and therefore carries no differential information.

A direction-agnostic differential posterior is obtained from the magnitude of the differential Bayes factor as *P* (diff | data) = |dBF|/(1 + |dBF|), and multiple testing across the shared-prey family is controlled by applying the same BFDR procedure used in the single-condition pipeline to these differential posteriors. Effect size is summarised by the difference in per-condition mean log_2_ fold change, Δ log_2_ FC = log_2_ FC_*A*_ − log_2_ FC_*B*_; by default each condition’s fold changes are *z*-scored within condition before differencing, so that protocol-level scale differences do not contaminate the contrast. This per-condition standardisation complements the bait-anchoring step (Equation 22), which equalises documented bait-abundance offsets between conditions by subtracting a single per-condition constant from every prey’s enrichment, leaving within-condition prey structure intact.

Each shared prey is then assigned to one of four states — *gained* (stronger in *A*), *reduced* (stronger in *B*), *unchanged*, or *bothnegative* — using per-condition interactor status (each condition’s BFDR against a threshold), the sign of Δ log_2_ FC, and significance of the differential BFDR proteins detected in only one condition are reported separately as condition-*A* - or condition-*B*-specific. Classification uncertainty is propagated rather than discarded: alongside the direct differential posterior error probability (PEP), a class-conditional posterior error probability is computed for each of the four states under an explicit conditional-independence approximation *P* (*s*_*A*_, *s*_*B*_ | data) ≈ *P* (*s*_*A*_ | data) ⋅ *P* (*s*_*B*_ | data), combining the per-condition interaction posteriors (Platt-calibrated where available) with logistic gates on Δ log_2_ FC. Finally, each protein receives a Bayesian-decision-theoretic call: under a default asymmetric 4 × 4 loss matrix that penalises direction flips most heavily and missed hits more than over-claims, the action minimising expected posterior loss is reported together with its expected risk, surfacing a ranked list of validation candidates whose differential call is both confident and decision-relevant.

The contrast generalises to *k* ≥ 3 conditions through all-pairs or one-versus-reference pairwise comparisons, with cross-pair multiple testing controlled by Benjamini– Hochberg (default), Bonferroni, or Holm correction over the flattened protein-by-contrast family, and an omnibus heterogeneity test — a closed-form Gaussian Bayes factor contrasting a shared-mean null against free per-condition means, with an empirical-Bayes pooled prior — flagging proteins whose enrichment differs across any condition. Each comparison emits a per-protein results table (Bayes factors, posteriors, BFDR, Δ log_2_ FC, classification, decision call) and a standard set of diagnostics — volcano, per-evidence, scatter, MA, and classification-count plots —coloured by interaction class.

#### Structural docking of high-confidence interactors

BayesInteractomics ships an optional, off-by-default structural-docking module that supplies an orthogonal, sequence-and-structure-based check on the most confident calls. The module is a request generator and result importer rather than a bundled docking engine: it does not run structure prediction itself. When docking is enabled, the package selects high-confidence interactors, emits AlphaFold Server^52^ batch-prediction request files for each bait–prey pair, and — after the user has run those predictions externally and returned the result archives –parses the predicted complexes and folds a calibrated structural Bayes factor into the posterior. Because prediction is user-mediated (the public AlphaFold Server exposes no programmatic interface and caps usage at roughly thirty jobs per day under a per-job token budget), the module organises requests into daily batches of at most thirty jobs, skips pairs whose combined bait-and-prey length exceeds the token budget, and deduplicates against a persistent on-disk cache keyed on the canonical (sorted) pair identity so that no pair is ever predicted twice across sessions.

Candidates are drawn from the final results table by requiring a posterior probability at or above a threshold (default 0.8) and a PEP at or below a threshold (default 0.01), retaining only detected proteins and, where mixture diagnostics were run, those flagged as well-behaved. Surviving candidates are ranked by a composite of the model posterior and a per-protein structural dockability score derived from AlphaFold-DB monomer confidence (mean predicted local-distance difference test and predicted dis-ordered fraction), prioritising the proteins for which a complex prediction is most informative; at most one hundred pairs are emitted by default. Each request docks the bait against one prey as a two-chain prediction with five random seeds; the bait sequence is supplied directly or auto-fetched from UniProt and is reused across all pairs. Alongside the per-pair and combined-batch request files, the module writes a machine-readable manifest, skip lists for cached and over-budget pairs, and a step-by-step upload guide.

Returned predictions are scored per pair from the five seeds. The interface confidence is summarised primarily by an AlphaFold3-calibrated composite of interface predicted local-distance difference test, interface predicted aligned error, predicted template-modelling score and interface predicted template-modelling score (the C2Qscore of^9^), with a per-residue pDockQ logistic^5354^ and an interface predicted-template-modelling-score step function available as lower tiers when the full per-residue prediction data are unavailable. The chosen interface score is mapped to a structural Bayes factor by a logistic calibration anchored to published precision points, and is passed through conservative quality gates that neutralise it (Bayes factor returned to one) for predictions dominated by disorder, low monomer confidence, weak inter-chain spatial proximity, or high seed-to-seed disagreement — the regimes in which interface confidence scores are known to be unreliable for transient and intrinsically disordered interactions. The structural evidence is then combined with the existing MS posterior by a two-stage sequential Bayesian update in log-odds space,

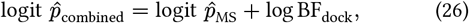

which is justified by the conditional independence of the two evidence channels: the structural score depends on predicted surface geometry and chemistry, whereas the MS observables (abundance ratios, cross-run correlation, detection counts) depend on co-elution in the affinity capture, so the two share no measurement pathway. Pairs without a usable prediction take BF_dock_ = 1 and are left unchanged, and the global BFDR is recomputed from the updated posteriors. The augmented table reports the original and combined posteriors side by side together with the raw structural scores, and a dedicated tab in the interactive report visualises the update, so the structural contribution is always auditable and never silently over-rides the MS evidence.

Because this update is anchored to a structural prediction of direct physical contact rather than to membership in a curated complex, it is independent of the reference databases used elsewhere for benchmarking. It therefore doubles as a reference-independent arbiter for high-confidence calls absent from curated interaction databases, distinguishing candidate direct binders from co-purifying indirect partners on evidence that no curated-recall ceiling can supply. We document the capability here as an implemented, optional refinement layer; a quantitative structural-validation study is reported in the companion application papers rather than claimed as a method-level result here.

### Validation studies

#### Synthetic benchmark

To compare scoring tools where the ground-truth interactome is known, we generated synthetic spike-in AP-MS datasets with a dual-branch generative model: prey spectral counts are drawn from a negative-binomial likelihood and a secondary log_2_-LFQ intensity from a Normal emission of the shared latent abundance, so that the count branch does not share the Gaussian family BayesInteractomics assumes for its LFQ evidence — a single-family generator would make the benchmark tau-tological. True interactors carry a graded per-interactor enrichment rather than a constant shift, populating the full range of effect sizes. The generator’s baseline count level, dispersion, and enrichment scale were calibrated so that the per-prey discriminability of each branch matches that measured on a real human deposit (OpenCell, PXD024909): on real data the LFQ contrast is more informative than spectral counts, and the generator is tuned to reproduce that ordering rather than the inverted, unrealistically clean count branch that naïve simulators produce which otherwise hands the count-based comparators an artificial advantage. The calibration targets — the single-feature and bait–control-contrast discriminability of each branch measured on the real deposit — were fixed before scoring and independently of any tool; the resulting match, and the inverted branch ordering produced by a naïve simulator, are reported in Extended Data Table 1. All stochastic steps are seeded, and the benchmark is run over *B* = 7 independently seeded batches.

Each batch is scored by all four tools through the same pipeline used throughout — SAINTexpress, CompPASS and MiST on spectral counts, BayesInteractomics on the LFQ branch; because the synthetic identifiers carry no STRING or structural annotation, BayesInteractomics runs copula-only, without its deep-learning prior, so its reported performance is a conservative floor. For each tool we compute the area under the ROC and precision–recall curves against the known labels; differences between BayesInteractomics and each comparator are summarised across the *B* batches by paired 95% boot-strap confidence intervals and a Wilcoxon signed-rank test, and reported as effect sizes with intervals rather than *p*-values alone.

## Data availability

The raw mass spectrometry data re-analysed in this work are publicly available through the PRIDE repository under the following accessions: Greco et al. 202210, PXD025510; Sap et al. 202112, PXD024254; and Hein et al. 2015, PXD002815. The HAP40 dataset is deposited with the companion study8 and has no standalone accession. The synthetic benchmark data and the comparator/validation pipeline are archived together in a single Zenodo record as a self-contained, version-pinned reproducibility package (zenodo.21217041). Per-figure Source Data files (one `.xlsx` per main-text statistical figure) accompany the manuscript.

## Code availability

The Julia package developed for this work, BayesInteractomics, is open-source under the MIT licence and available on GitHub (https://github.com/ma-seefelder/BayesInteractomics). The repository includes the complete source code, documentation, the trained deep neural network and meta-learner models, and scripts to reproduce the analyses presented in this study. The exact source snapshot documented here (v1.2.1) is archived on Zenodo (zenodo.21217041), together with a self-contained reproducibility package that regenerates every main-text and supplementary figure from its committed intermediate data and includes the per-figure Source Data.

## Acknowledgements

This study was supported by the European Huntington Disease Network (EHDN) via their Lesley Jones Seed Fund Program (project number: 1245). We thank Fabrice Klein and Stefan Kochanek for support and helpful discussions. Furthermore, we thank Todd Greco for providing their AP-MS data on the HTT interactome for reanalysis^11^.

## Author contributions

M.S. conceptualised the study, developed the methodology, designed and implemented the software, performed the data analysis and validation, wrote the original draft, and reviewed and approved the final manuscript.

## Additional Information

**Correspondence and requests for materials** should be addressed to Manuel Seefelder.

## Extended Data

**Extended Data Fig. 1.**
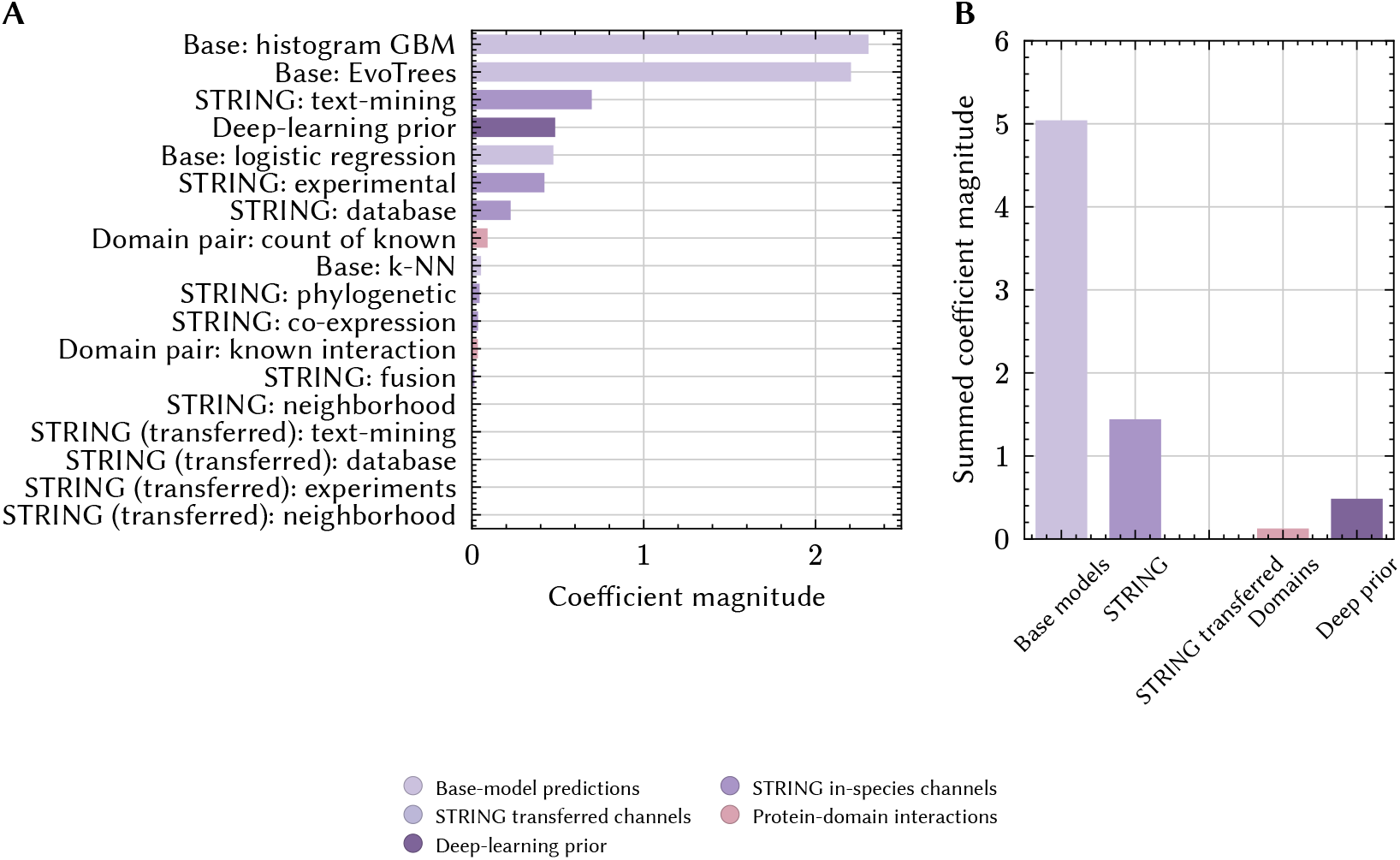
Blender coefficients of the stacked meta-learner. Absolute coefficient magnitudes of the L2-regularised logistic-regression blender on standardised inputs. (A) Per-input coefficient for each of the 18 inputs the blender sees: the 14 raw features plus the four out-of-fold base-learner predictions produced by the stack’s internal cross-validation. (B) The same coefficient magnitudes aggregated by feature family.

**Extended Data Fig. 2.**
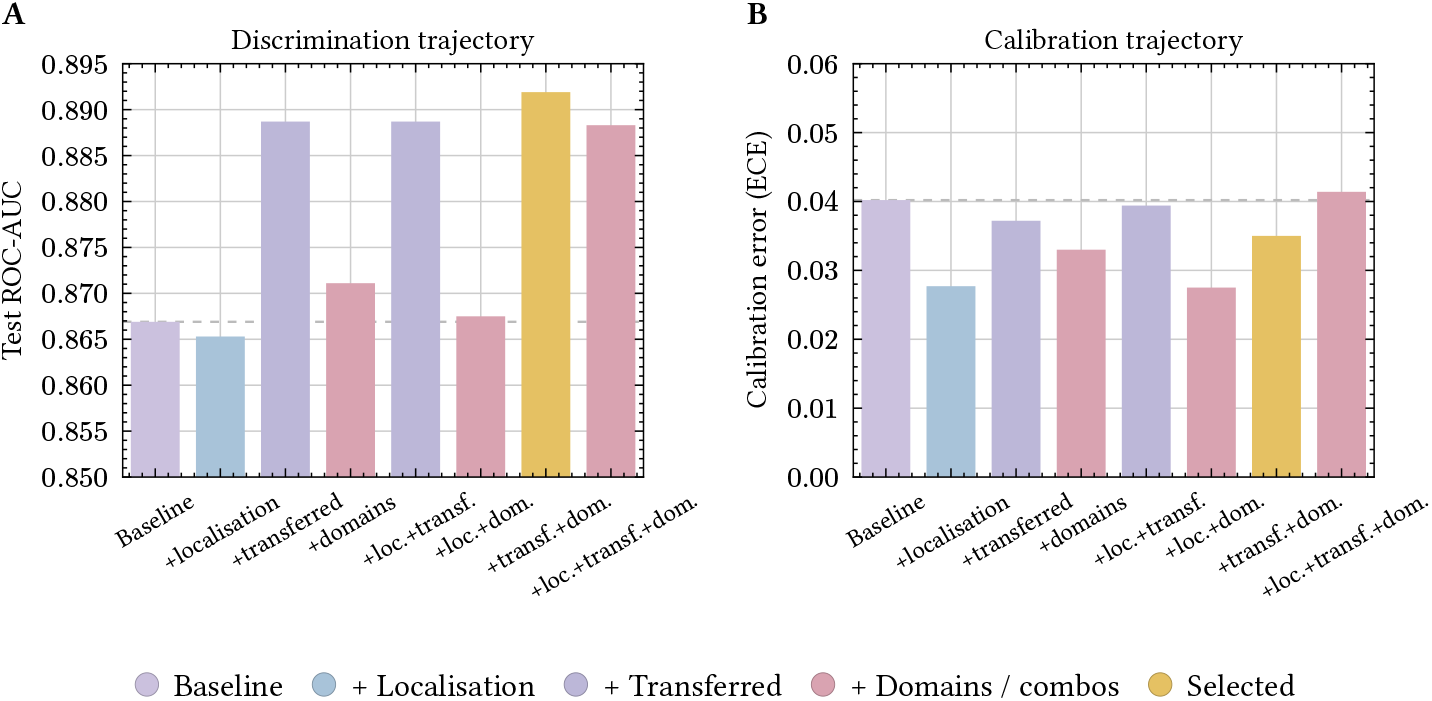
Discrimination and calibration as feature families are added to the meta-learner blender. Starting from a baseline of seven in-species STRING channels plus the DNN prior (and the four base-learner out-of-fold predictions), subcellular-localisation features, cross-species transferred STRING channels, Pfam protein-domain features, and Gene Ontology semantic-similarity features are added in all single and multi-family combinations. (A) Test-set ROC-AUC. (B) Test-set calibrated ECE. The selected configuration is transferred + domains on top of the baseline, which lifts ROC-AUC by +0.017 over baseline and lowers ECE by ≈ 25%. Dashed lines mark the baseline value.

**Extended Data Fig. 3.**
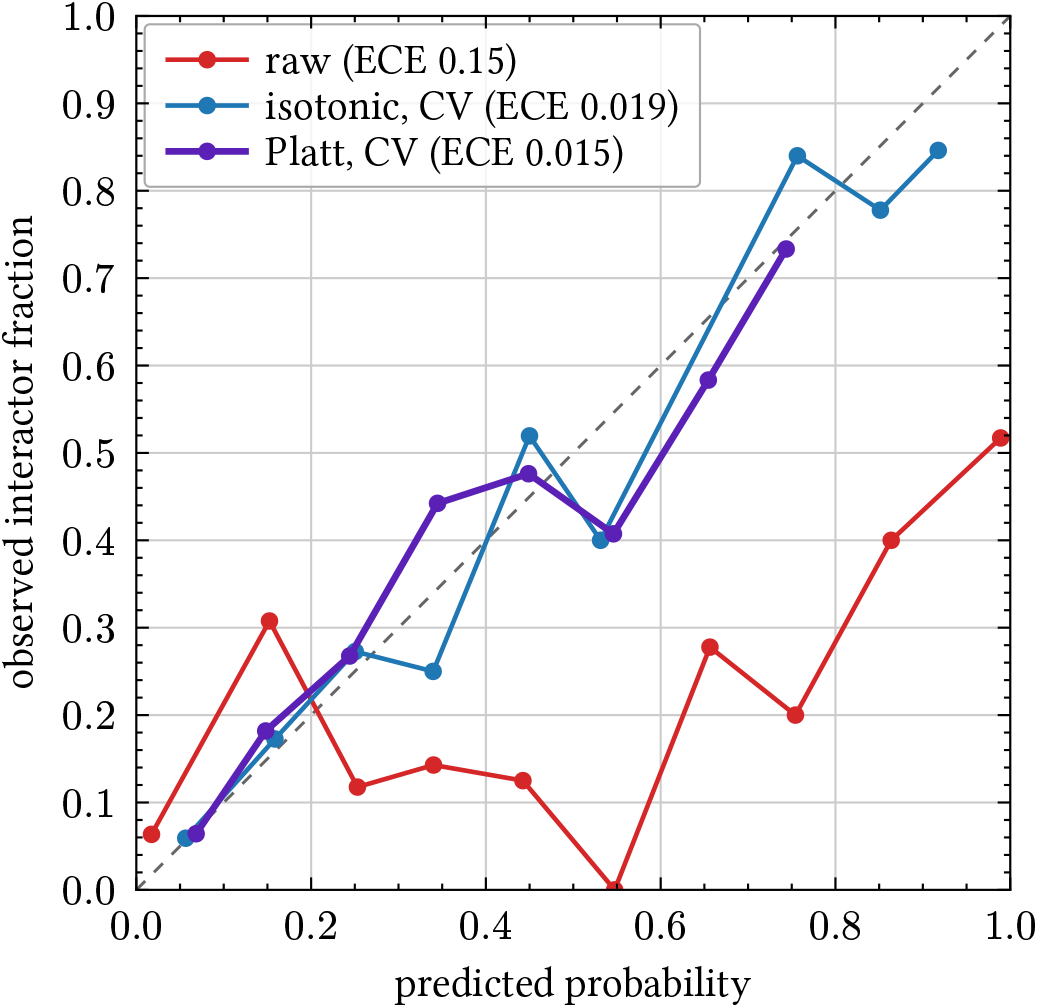
Posterior calibration on synthetic complete ground truth. Reliability diagram of the BayesInteractomics combined posterior on a graded-signal synthetic substrate (*n* = 1500 candidate proteins, 16.3% true interactors) for which ground truth is complete. The *raw* posterior lies below the diagonal and is over-confident (ECE 0.15, MCE 0.55). A held-out, five-fold cross-validated Platt recalibration fit *against the known labels* returns the curve to the diagonal (ECE 0.015, MCE 0.14), and a cross-validated isotonic fit reaches a comparable value (ECE 0.019). Because the recalibration is monotone, the area under the ROC curve is invariant (0.83). In routine use the pipeline’s recalibration is fit on the model’s own simulated ground truth rather than on external labels.

**Extended Data Table 1:** Calibration of the synthetic generator to real per-prey discriminability. The spike-in generator’s per-prey separability was tuned to an external target — the median single-feature and bait–control-contrast AP-MS discriminability measured across a real human deposit (OpenCell, PXD024909) — fixed from real data before scoring and independently of any tool. A naïve simulator (constant enrichment, conventional count parameters) makes both branches almost perfectly separable and the real branch ordering; the calibrated generator matches the real count branch and both bait–control-contrast targets and restores the real ordering, in which the LFQ contrast is more informative than spectral counts. The calibrated LFQ single-feature separability (0.824) runs above the real median (0.667), while the count branch and both contrast metrics match real data. Values are replicate-averaged AUROC on a representative seed; the calibrated generator is the substrate used in Extended Data Fig. 4.

| 1565 | Per-prey discriminability (median AUROC) | Real OpenCell (target) | Naïve simulator | Calibrated generator |
| --- | --- | --- | --- | --- |
| 1566 | Count branch, single-feature | 0.755 | 0.975 | 0.758 |
| 1567 | LFQ branch, single-feature | 0.667 | 0.906 | 0.824 |
| 1568 | Count branch, bait-control contrast | 0.641 | 0.984 | 0.629 |
| 1569 | LFQ branch, bait-control contrast | 0.726 | 0.877 | 0.764 |
| 1570 | Branch ordering (contrast) | LFQ > count | count > LFQ | LFQ > count |

**Extended Data Fig. 4.**
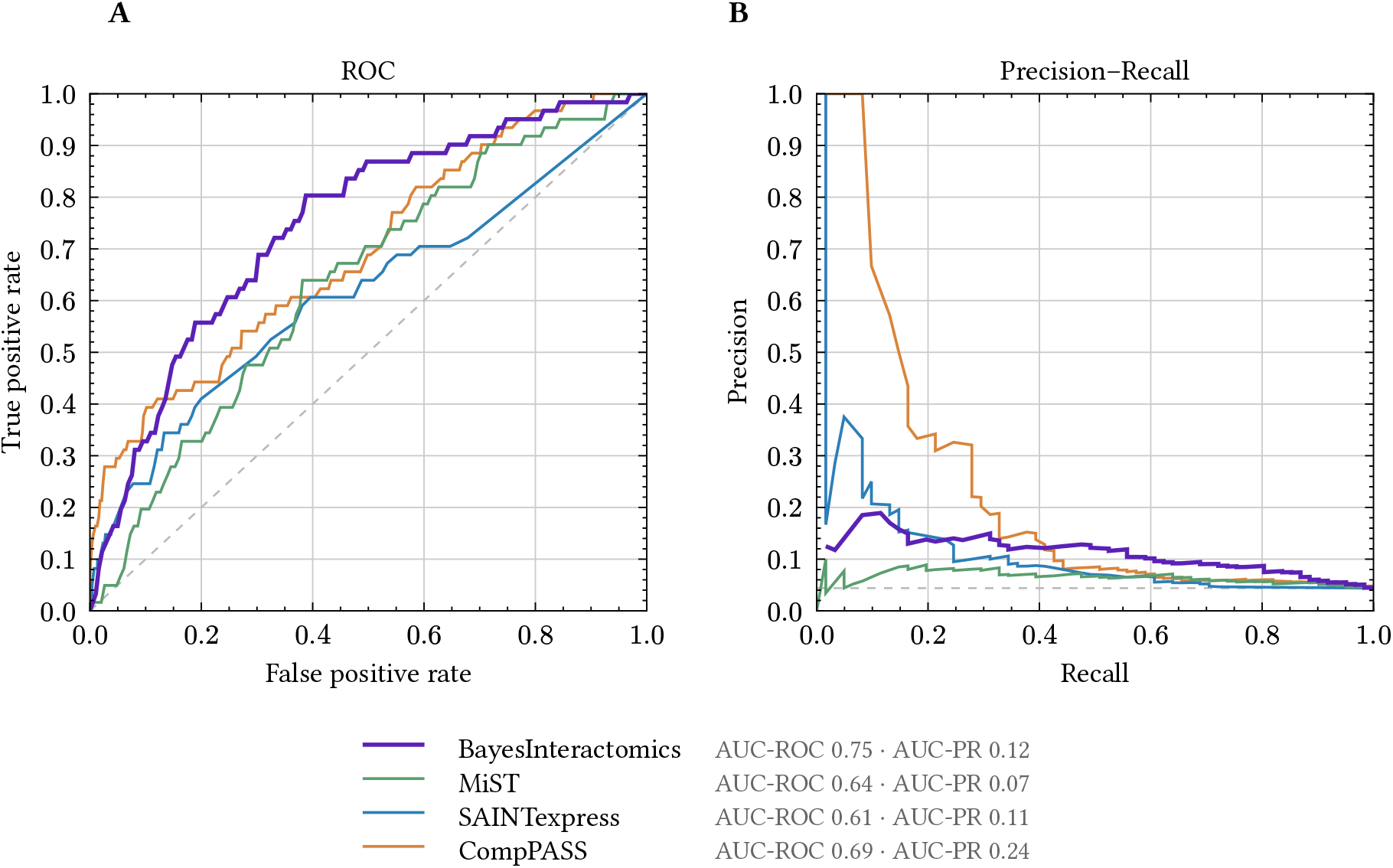
Four-tool comparison on a realistically-calibrated synthetic substrate. All four scorers were run on a spike-in synthetic AP-MS substrate with complete known truth (61 true interactors among 1,389 candidates, positive rate 4.4%), the generator’s per-prey discriminability calibrated to real AP-MS data so that no tool receives an artificially easy substrate (Extended Data Table 1). (A) Pooled ROC and (B) precision–recall curves for the representative AUROC-median batch of *B* = 7 seeds; dashed lines mark chance (the diagonal and the synthetic positive rate, respectively). On this substrate the count-based comparators score 0.61–0.69 AUROC, against the 0.97–0.99 they reach on idealised simulators. BayesInteractomics was run copula-only, without its deep-learning prior. Paired across the *B* = 7 seed batches (BayesInteractomics − comparator), ΔAUROC is +0.134 [0.100, 0.156] versus SAINTexpress, +0.068 [0.034, 0.099] versus CompPASS and +0.128 [0.091, 0.155] versus MiST (all CIs > 0; Wilcoxon signed-rank *p* = 0.016); on precision– recall BayesInteractomics leads SAINTexpress (+0.083) and MiST (+0.112) and trails CompPASS (−0.040). Discrimination and the calibratability of the synthetic posterior are characterised separately (Extended Data Fig. 3).

**Extended Data Fig. 5.**
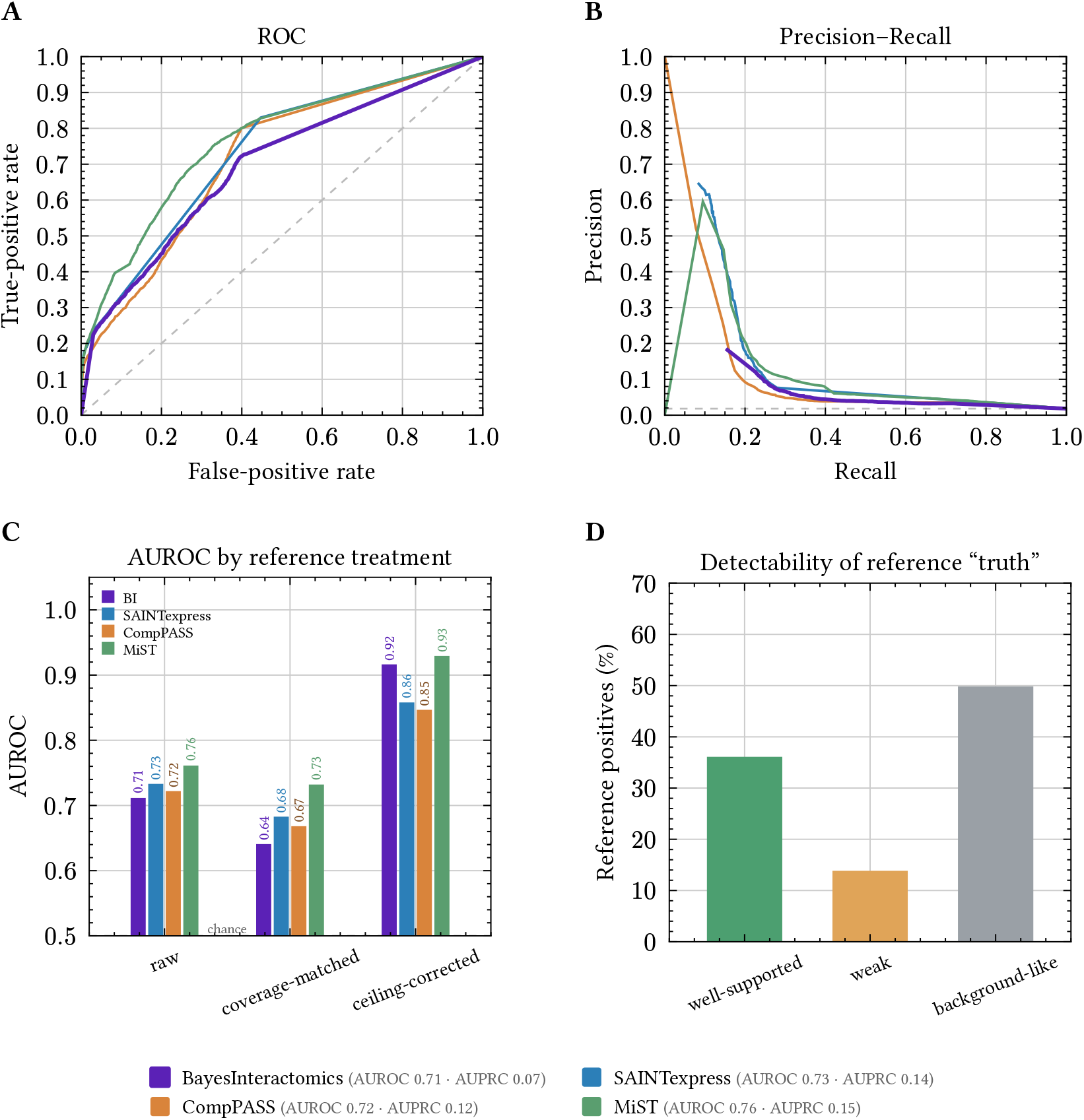
Real-data benchmark on a second, independent human dataset (Hein 2015). All four scorers were run on the Hein 2015 green fluorescent protein (GFP)-overexpression AP-MS panel (PXD002815; 75 baits, 2,651 reference positives) and scored against the same external CORUM ∪ IntAct reference as Figure 2; BayesInteractomics and SAINTexpress run per bait, CompPASS and MiST whole-panel. (A) Pooled ROC and (B) precision–recall curves onthe raw scores; dashed lines mark chance. (C) AUROC under the same three reference treatments as Figure 2 — *raw, coverage-matched*, and *ceiling-corrected* (positives restricted to the well-supported tier). (D) Detectability of the curated “truth”: 36% of reference positives are well-supported, 14% weak and 50% background-like — the last carry no usable signal and are forced false negatives for every method. On the raw and coverage-matched views BayesInteractomics sits mid-panel; on the ceiling-corrected view it ranks second (ROC-AUC 0.92, behind MiST 0.93 and ahead of SAINTexpress 0.86 and CompPASS 0.85), its posterior tracking measured per-prey enrichment second best (Spearman *ρ* = 0.54). Open *world* / *incomplete reference:* undetected or uncurated CORUM/IntAct prey are ranked last and the reference is partly circular, so these metrics are indicative and do not constitute a complete accuracy ranking. *Overexpression / scale context*: Hein 2015 uses GFP-tagged overexpression constructs (not endogenous), a distinct experimental regime from OpenCell, so AUROC/AUPRC values are not directly comparable across datasets.

**Extended Data Fig. 6.**
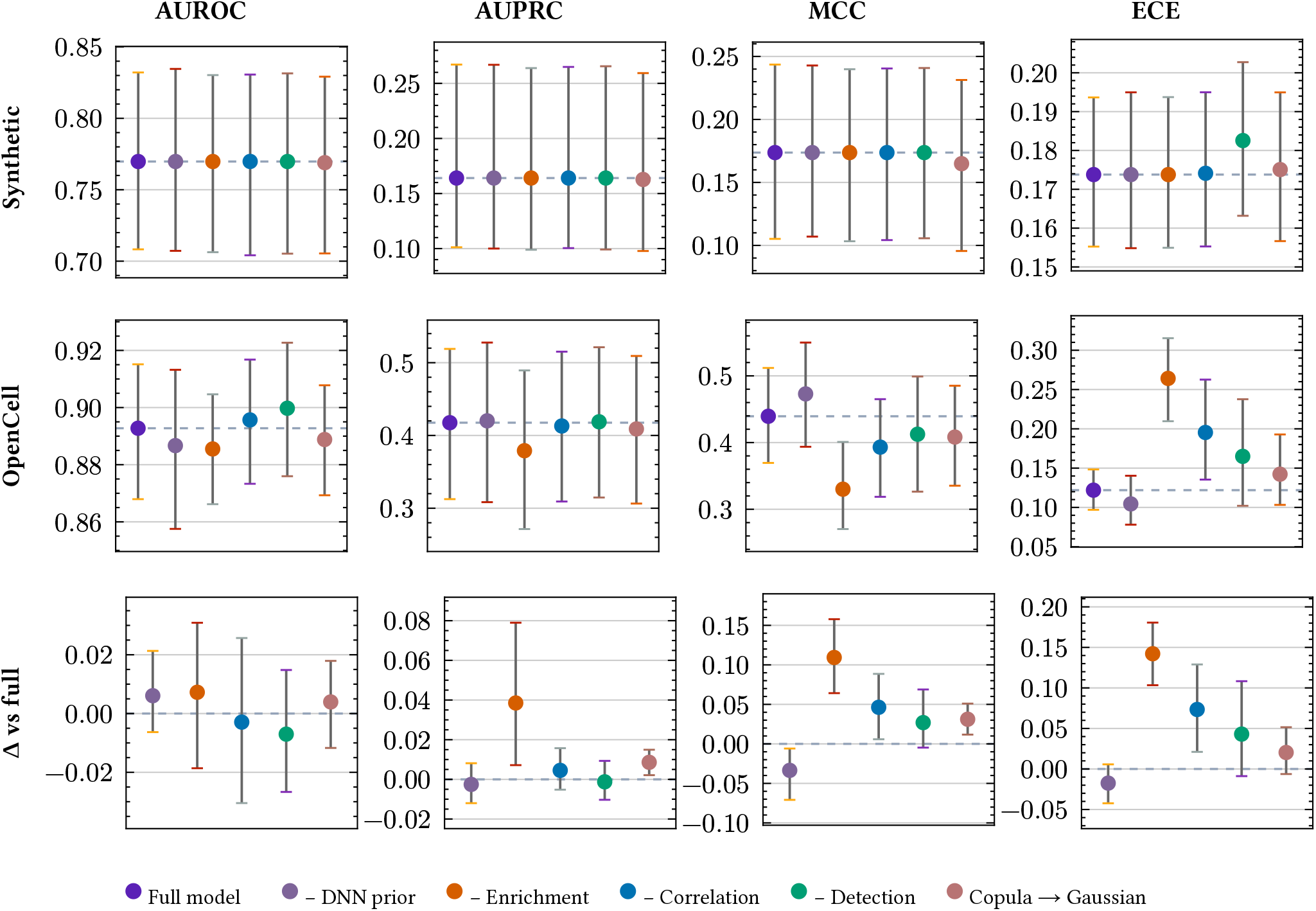
Architectural ablation of BayesInteractomics components. Each component is dropped or swapped in turn — the DNN prior, each of the three evidence streams (enrichment, correlation, detection), and the copula-family selection (BIC-auto → forced Gaussian) — each a full model re-fit via the native v1.2.1 CONFIG knobs (not a parameter tweak). Four metrics: AUROC and AUPRC (ranking quality), MCC at threshold 0.5 (binary-call quality) and ECE (calibration;*lower* is better); points = value, whiskers = bootstrap 95% CI. Top row: synthetic fair-generator substrate (complete ground truth, bootstrap over preys; dashed line = full model). Middle row: OpenCell, *ceiling-corrected* (positives restricted to the well-supported tier and coverage-matched to the full-model scored set — the same honest treatment as Figure 2; bootstrap over 12 baits; dashed line = full model). Bottom row: the *paired* contribution of each component on OpenCell — Δ versus the full model computed per bait on the same baits, so the between-bait heterogeneity that widens the absolute intervals cancels (paired bootstrap over 12 baits; positive = the component helps; dashed line = no effect; a CI clear of 0 marks a resolved contribution). On the synthetic substrate the three streams are mutually redundant (every component sits on the full-model line). On real data, ranking (AUROC) is robust to every single drop, whereas enrichment contributes resolvably to AUPRC (+0.039, [+0.007, +0.079]), MCC (+0.109, [+0.064, +0.158]) and calibration (ECE +0.142, [+0.103, +0.181]); correlation and the copula-family choice contribute resolvably to MCC and (correlation) to ECE, and the copula choice carries a small AUPRC effect (+0.009, [+0.002, +0.015]). The DNN prior is feature-limited on this dataset — its STRING and protein-domain look-ups are unavailable for these prey identifiers (matching the head-to-head run) — so dropping it is near-neutral to slightly beneficial here (see Table 1 for its meta-learner-level evaluation). The full variant reproduces the Figure 2 BayesInteractomics result. Source numbers in `figure_ablation_source_data.xlsx`.

